# Estimating the Selective Effect of Heterozygous Protein Truncating Variants from Human Exome Data

**DOI:** 10.1101/075523

**Authors:** Christopher A. Cassa, Donate Weghorn, Daniel J. Balick, Daniel M. Jordan, David Nusinow, Kaitlin E. Samocha, Anne O'Donnell Luria, Daniel G. MacArthur, Mark J. Daly, David R. Beier, Shamil R. Sunyaev

## Abstract

The dispensability of individual genes for viability has interested generations of geneticists. For some genes it is essential to maintain two functional chromosomal copies, while other genes may tolerate the loss of one or both copies. Exome sequence data from 60,706 individuals provide sufficient observations of rare protein truncating variants (PTVs) to make genome-wide estimates of selection against heterozygous loss of gene function. The cumulative frequency of rare deleterious PTVs is primarily determined by the balance between incoming mutations and purifying selection rather than genetic drift. This enables the estimation of the genome-wide distribution of selection coefficients for heterozygous PTVs and corresponding Bayesian estimates for individual genes. The strength of selection can help discriminate the severity, age of onset, and mode of inheritance in Mendelian exome sequencing cases. We find that genes under the strongest selection are enriched in embryonic lethal mouse knockouts, putatively cell-essential genes inferred from human tumor cells, Mendelian disease genes, and regulators of transcription. Using an essentiality screen, we find a large set of genes under strong selection that are likely to have critical function but that have not yet been studied extensively.

---

The evolutionary cost of gene loss is a central question in genetics and has been investigated in model organisms and human cell lines^1–3^. In humans, the question of dispensability and haploinsufficiency of individual genes is intimately related to their causal role in genetic disease. However, estimates of the selection and dominance coefficients in humans have proved elusive as inference techniques used in other sexual organisms generally require cross-breeding over several generations.

The analysis of patterns of natural genetic variation in humans provides an alternative approach to estimating selection intensity and dispensability of individual genes. Despite substantial methodological progress in the ascertainment and analysis of population sequence data^4–8^, estimation of parameters of natural selection in humans has been complicated by genetic drift, complexities of human demographic history^4,7,9–13^ and the role of non-additive genetic variation^14–16^. Additionally, naturally occurring PTVs are infrequent in the population and as a result, datasets of even thousands of individuals are underpowered for the estimation of gene dispensability in humans.

The Exome Aggregation Consortium (ExAC) dataset now provides a sufficiently powered sample to assess the selection that constrains the number of gene-specific PTVs in the general population^17^. We restrict our analysis to PTVs predicted to be consequential^18^, which allows us to assume that all PTVs within a gene likely incur the same selective disadvantage. We can then treat each gene as a bi-allelic locus with a functional state and a loss-of-function state. In each gene, the cumulative frequency of rare deleterious PTVs (the sum of PTV allele frequencies throughout the gene) is then primarily determined by the balance between incoming mutations and selection rather than through reassortment of alleles by stochastic drift. This makes our estimates robust to drift, population structure and historical changes in population size, which we evaluate analytically and with simulations (**Methods and Supplementary Figure 1**).

**Figure 1:**
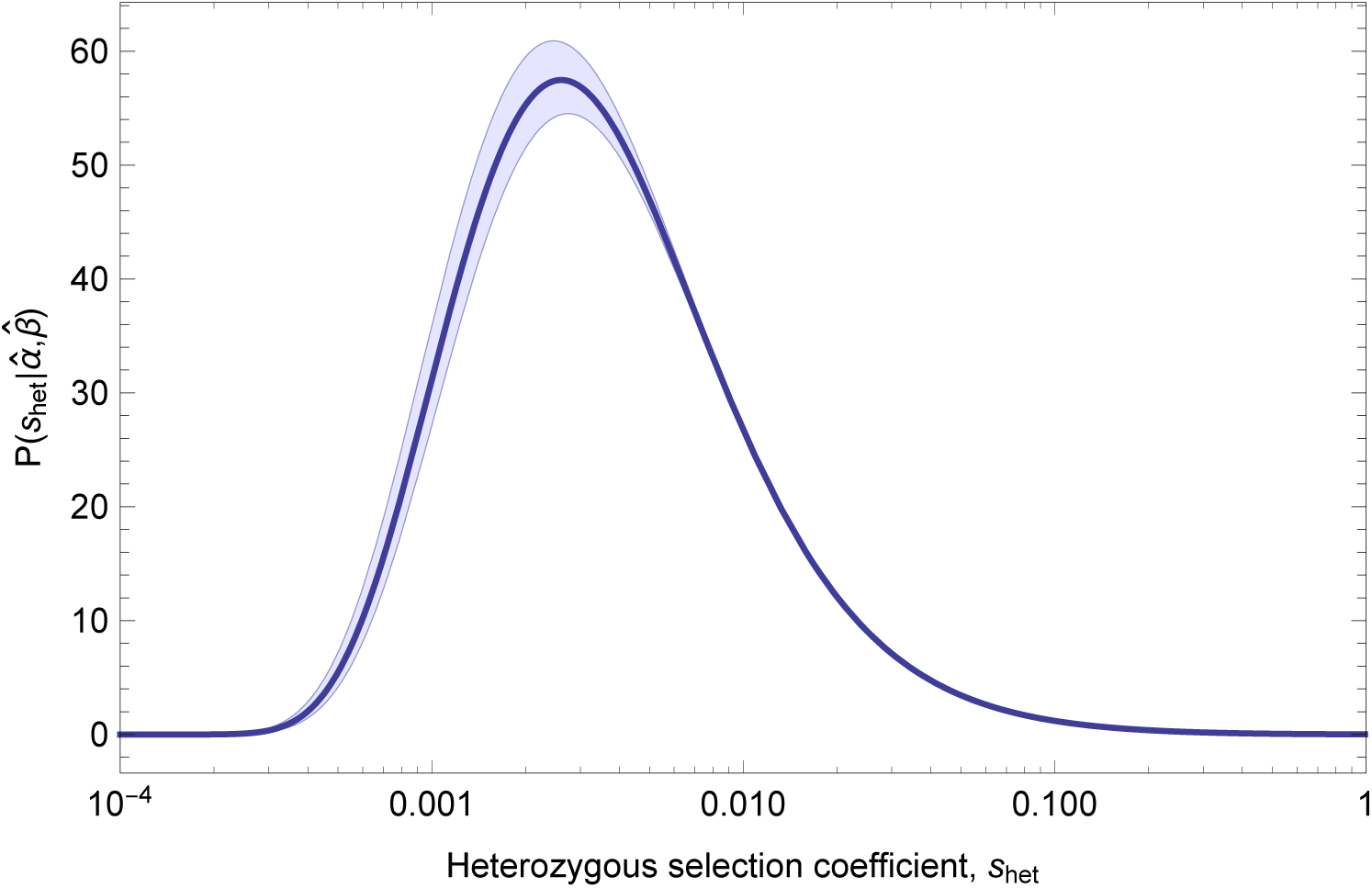
Inferred distribution of fitness effects for heterozygous loss of gene function. Estimates of parameters 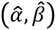from maximum likelihood fit to the observed distribution of PTV counts across 15,998 genes in terciles of mutation rate, assuming *S*_het_~IG (α, β). Shaded areas show 95% CI obtained from 100 bootstrapping replicates, intended to quantify the influence of sampling noise in the data set on parameter inference, with fixed estimates of local mutation rate.

Using population frequency data from 60,706 jointly-called exomes from individuals without severe Mendelian disorders, we estimate both the overall distribution of gene-based fitness effects and individual gene fitness cost in heterozygotes. Given gene-specific estimates of the *de novo* mutation rate^19,20^, the observed number of PTV alleles throughout each gene, and the number of chromosomes sampled, we estimate the distribution of the genome-wide selective effect of PTVs on heterozygote carriers, *S*_het_. We parameterize the distribution of selective effects using an inverse Gaussian, which is fit using maximum likelihood (**Figure 1**). We then estimate the selection coefficient for each gene using the posterior probability for *S*_het_ given gene-specific values of the observed number of PTVs, the number of chromosomes sampled and the estimated mutation rate (**Supplementary Table 1**).

Although the distribution is broad, suggesting that the effect of losing one copy of a gene is variable, the mode of the distribution corresponds to a fitness loss of about 0.5% (*S*_het_ = 0.005). Despite the large sample size, resolution to distinguish between very high selective effects is still limited. There are 2,984 genes with *S*_het_ > 0.1, a result concordant with previous estimates of loss of function intolerance derived from population data^17^. Even though some genes are heavily depleted of PTVs in ExAC as compared with mutational expectation, these values suggest that heterozygote PTVs in many genes are not necessarily responsible for observable, severe clinical consequences.

Unsurprisingly however, genes known to be involved in rare Mendelian diseases have higher *S*_het_ values. Among them, genes annotated exclusively as autosomal dominant (AD, N=867) have significantly higher *S*_het_ values than those annotated as autosomal recessive (AR, N=1,482)^21^ [Mann-Whitney p-value 3.14×10^−64^] (**Figure 2[a,b]**). This suggests that it may be possible to prioritize candidate disease genes identified in clinical exome sequencing analysis using the observed mode of inheritance and *S*_het_ value.

**Figure 2:**
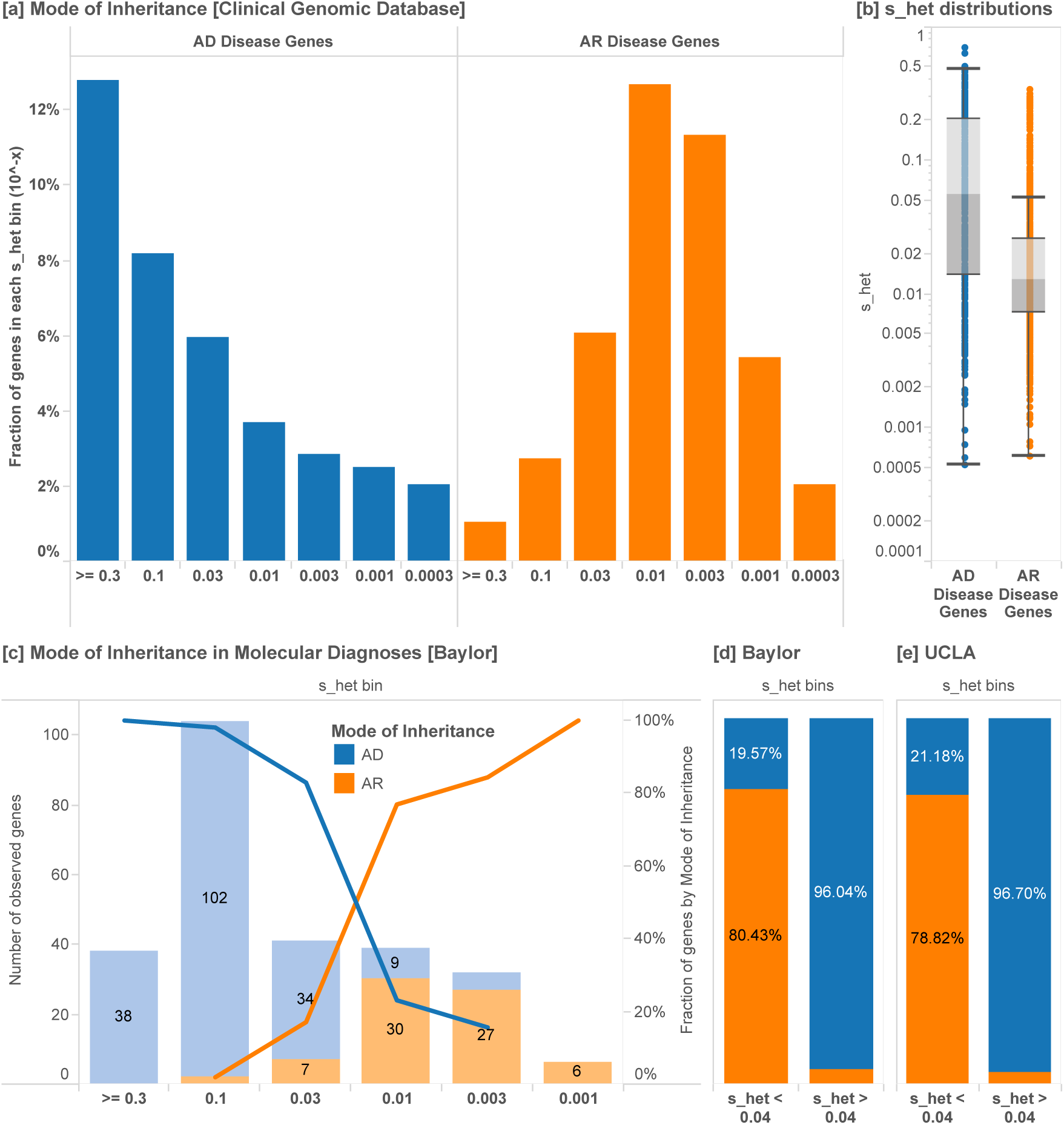
Separation of disease genes and clinical cases by mode of inheritance. [a] The distribution of genes associated with exclusively autosomal dominant (AD, N=867) disorders versus autosomal recessive (AR, N=1,482) disorders as annotated by the Clinical Genomics Database (CGD). Logarithmic bins are ordered from greatest to smallest *S*_het_ values. [b] Overall, AD genes have significantly higher *S*_het_ values than AR genes [Mann-Whitney p-value 3.14×10^−64^]. [c] Similarly, in solved Mendelian clinical exome sequencing cases (Baylor)^22^, *S*_het_ values can help discriminate between AR and AD disease genes, as annotated by clinical geneticists. [d] A *S*_het_ value of 0.04 can be used as a simple classification threshold for AD genes with a PPV of 96%. [e] This finding is replicated in a separately ascertained sample from UCLA. Box plots range from 25^th^–75^th^ percentile values and whiskers include 1.5 times the interquartile range.

In a set of 504 clinical exome cases that resulted in a Mendelian diagnosis^22^, we find a similar enrichment of cases by MOI and selection value (**Figure 2[c]**). We find that 90.4% of novel, dominant variants are associated with heterozygous fitness loss greater than 0.04 (**Figure 2[d]**). Among disease variants, a cutoff of *S*_het_ > 0.04 provides a 96% positive predictive value for discriminating between AD and AR modes of inheritance.

To test the generalizable utility of *S*_het_ values in prioritizing candidate genes in Mendelian sequencing studies, we compared the overall prevalence of genes with *S*_het_ > 0.04 to the corresponding fraction in an independently ascertained dataset of new dominant Mendelian diagnoses (**Figure 2[e]**)^23^. This analysis suggests that restricting to genes with *S*_het_ > 0.04 would provide a three-fold reduction of candidate variants, given the overall distribution of *S*_het_ values. Thus, initial effort in clinical cases can be focused on just a few genes for functional validation, familial segregation studies, and patient matching. We summarize the classification accuracy for all possible thresholds (AUC 0.9312) and probabilities for the mode of inheritance in each gene, generated using the full set of clinical sequencing cases (**Supplementary Figure 2 and Supplementary Table 2**).

Beyond mode of inheritance, we find that *S*_het_ can help predict phenotypic severity, age of onset, penetrance, and the fraction of *de novo* variants in a set of high-confidence haploinsufficient disease genes (**Figure 3**). In broader sets of known disease genes, *S*_het_ estimates significantly correlate with the number of references in OMIM MorbidMap and the number of HGMD disease “DM” variants (**Supplementary Figure 3**).

**Figure 3:**
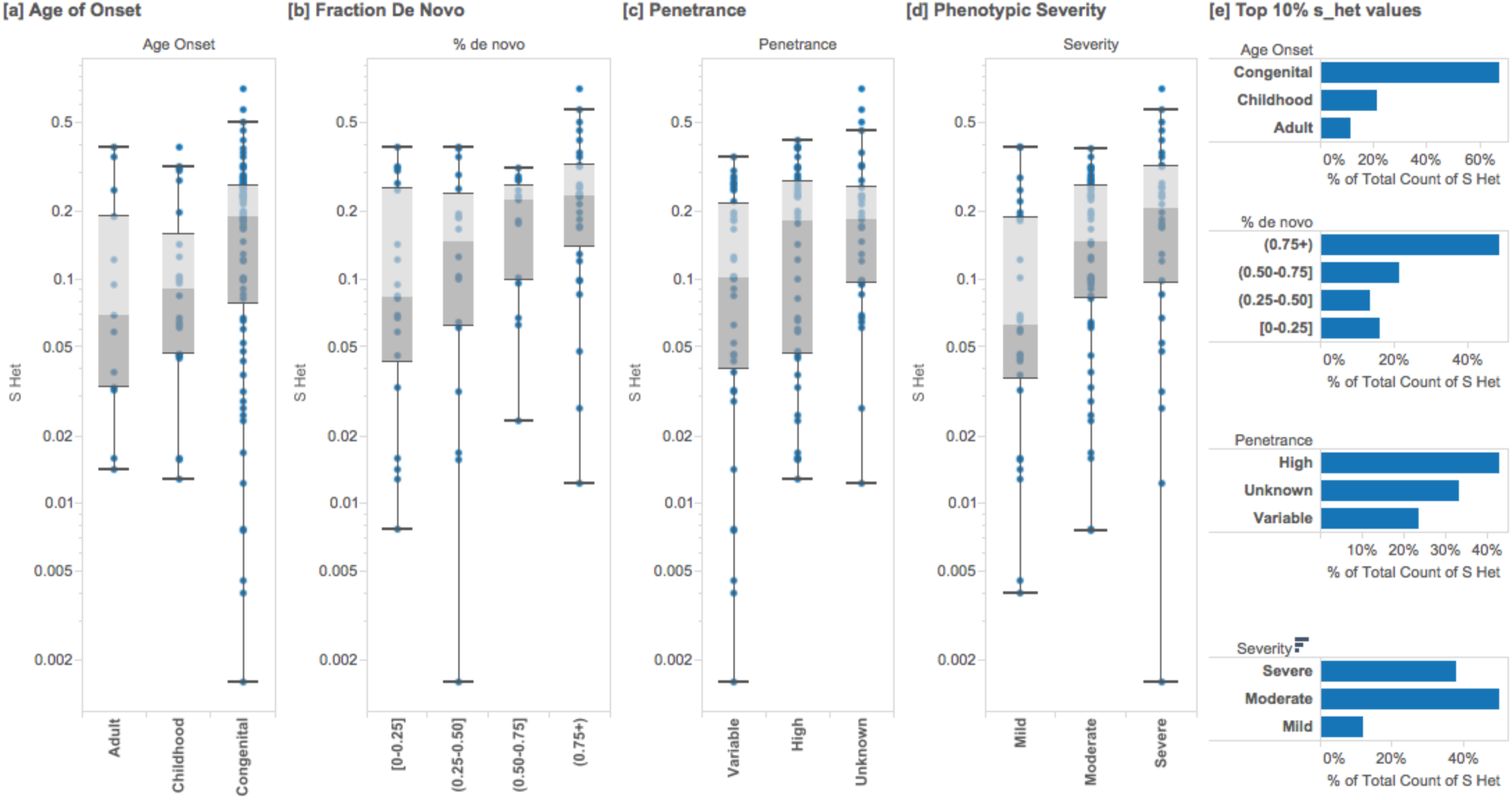
Enrichments of *S*_het_ in known haploinsufficient disease genes of high confidence (ClinGen Project). In (N=127) autosomal genes, we annotate the *S*_het_ scores of genes associated with each disease category and classification. Higher *S*_het_ values are associated with increased phenotypic severity (Mann-Whitney p-value 4.87×10^−3^), earlier age of onset (p=1.46 ×10^−2^), high or unspecified penetrance (p=1.79×10^−2^), and a larger fraction of *de novo* variants (p=8×10^−5^). Box plots range from 25^th^-75^th^ percentile values and whiskers include 1.5 times the interquartile range.

Gene-specific fitness loss values allow us to plot the distribution of selective effects for different disorders. This provides information about the breadth and severity of selection associated with various disorder groups using both well-established genes (**Figure 4[a]**) and new findings from Mendelian exome cases (**Figure 4[b]**). Overall, genes involved in neurologic phenotypes and congenital heart disease appear to be under more intense selection when compared with other disorder groups, tolerated knockouts in a consanguineous cohort, or in all genes (**Figure 4[c,d]**)^24^. Interestingly, genes recessive for these disorders appear to have only partially recessive effects on fitness, so selection on heterozygotes is not negligible in these genes (**Figure 4)**.

**Figure 4:**
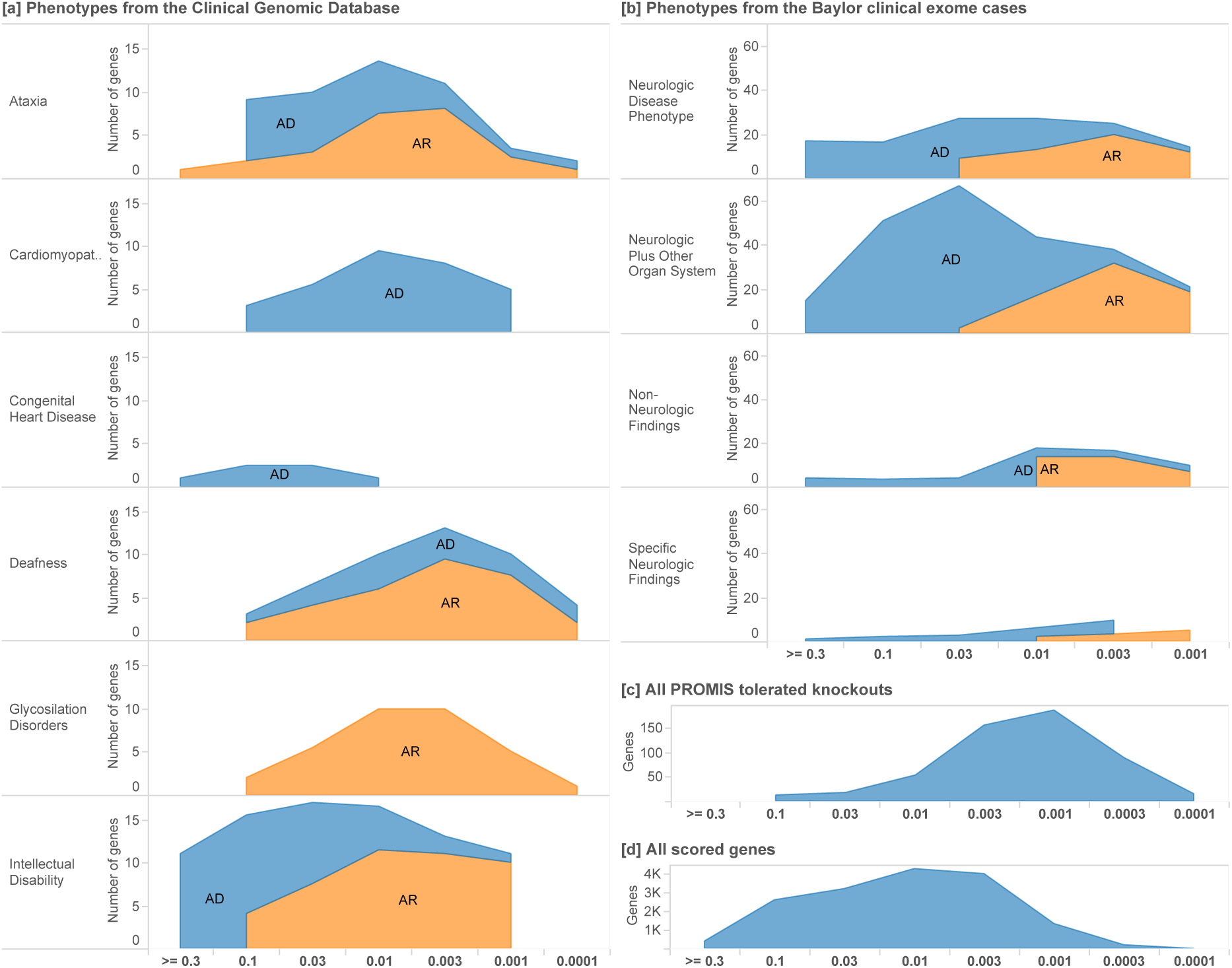
Distribution of *S*_het_ values for phenotypes in known disease genes and clinical cases. We plot the distribution of selective effects for different disorder groups, providing information about the breadth and severity of selection associated with each group. [a] We include known Mendelian disease genes (Clinical Genomic Database) annotated as either Autosomal Recessive or Autosomal Dominant and [b] clinical exome sequencing cases^22^. We contrast these with [c] all tolerated knockouts in a consanguineous cohort (PROMIS)^24^ and [d] the distribution of selective effects in all scored genes. Logarithmic bins are ordered from greatest to smallest *S*_het_ values.

In germline cancer predisposition, genes with higher selection values are enriched in individuals with cancer over those in ExAC (**Supplementary Figure 4**). This suggests that genes with low *S*_het_ values should not be prioritized in the prospective genetic screening for cancer predisposition. Consistent with previous studies^19^, we find that *de novo* mutations in patients with autism spectrum disorder are significantly enriched in genes with higher selective effects than those identified in controls (**Supplementary Figure 5 and Supplementary Table 3**).

**Figure 5:**
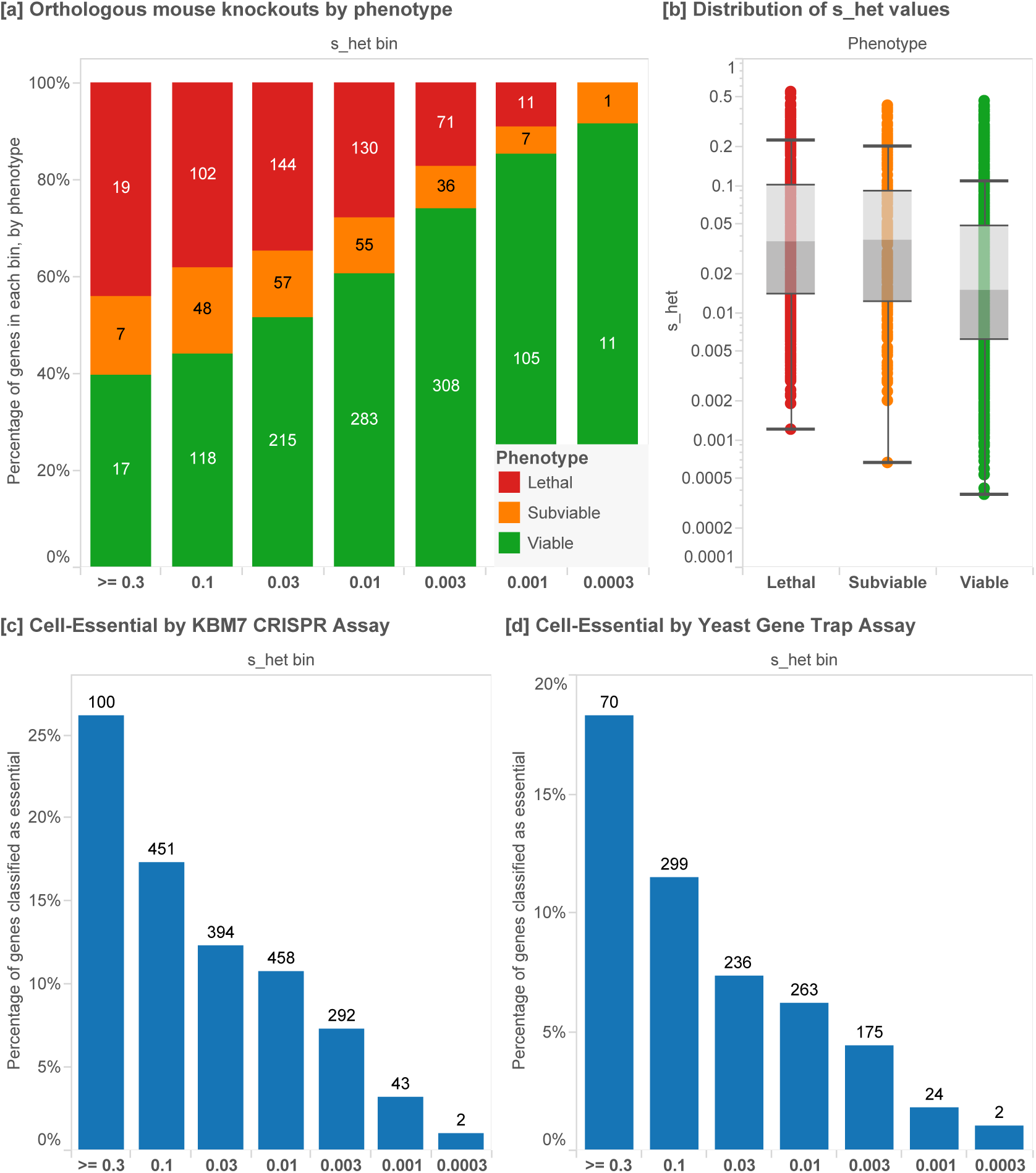
High-throughput screens of gene essentiality in mice and cell assays. [a] Proportion of orthologous mouse knockout genes by phenotype, from a neutrally-ascertained set of genes generated by the International Mouse Phenotyping Consortium (IMCP). Logarithmic bins are ordered from greatest to smallest *S*_het_ values. [b] ICMP mice are separated into viable (N=1,057), sub-viable (N=211) and lethal knockouts (N=477), and lethal knockouts have significantly higher *S*_het_ values than viable [Mann-Whitney p-value 2.95×10^−28^]. [c] Cell-essential genes as reported by Wang *et al.* from genome-wide KBM-7 tumor cell CRISPR assay (N=1,740) have significantly higher *S*_het_ values [p-value 5.13×10^−16^] and [d] as do genes that were characterized as essential in a gene trap assay (N= 1,081) [p-value = 4.90×10^−18^]. In the CRISPR assay, all genes with adjusted p-values < 0.05 and negative assay scores are included, and genes with gene trap scores < 0.4 or lower are included. Box plots range from 25^th^–75^th^ percentile values and whiskers include 1.5 times the interquartile range.

Next, we analyze *S*_het_ in the context of developmental and functional assays. In a large set of neutrally-ascertained mouse knockouts (N=2,179 genes)^25^, mice that are null mutant for orthologous genes with higher *S*_het_ estimates are enriched for embryonic lethality or sub-viability, while those with the lowest *S*_het_ estimates are depleted for embryonic lethality [Mann-Whitney p=2.95×10^−28^] (**Figure 5[a,b]**).

It is well known that mutations that are haploinsufficient in humans can often be well-tolerated when heterozygous in mice^26^. A classic example is *SHH*; heterozygous null mutations in this important developmental signaling gene result in holoprosencephaly^27^. Haploinsufficiency for other genes in this signaling pathway also results in developmental defects; e.g. *GLI3* (Pallister-Hall syndrome and Greig cephalopolysyndactyly syndrome)^28–30^ and *GLI2* (Holoprosencephaly 9)^31^. Interestingly, haploinsufficiency for these genes is tolerated in mouse models; mice heterozygous for null variation in the *SHH* signaling pathway are phenotypically normal, while homozygous mutant mice have defects that recapitulate features of the human syndrome^32–34^. This extends to many other human developmental disorders, enabling the experimental characterization of the molecular consequences of these mutations. Thus, it is notable that mice that are homozygous for null mutations in orthologous genes with higher *S*_het_ values are enriched for lethality.

High-throughput genetic analysis of cell-essentiality provides an orthogonal dataset for comparison with our estimates of *S*_het_. In genes that are predicted to be essential for human cell proliferation using CRISPR-based inactivation (**Figure 5[c]**) and gene trap inactivation assays^3^(**Figure 5[d]**), we find that putatively essential genes are heavily enriched in genes with high *S*_het_ values [p-values 5.13×10^−16^, 4.90×10^−18^, respectively].

Key developmental pathways are dramatically enriched in genes with high *S*_het_ values (**Figure 6[a]**). We also find a significant positive correlation between the number of protein-protein interactions for each gene and its *S*_het_ value (**Figure 6[b,c]**), identified from high-throughput mass spectroscopy data. In the context of molecular and cellular function, a set of genes with very high estimated selective effects (*S*_het_ > 0.15, 2,072 genes) is statistically enriched in biological process categories “transcription regulation” (Bonferroni corrected p=1.8×10^−39^), “transcription” (7.5×10^−36^), and “negative regulators of biosynthetic processes” (see **Supplementary Material**)^35^. Consistent with this finding, nucleus was the most enriched cellular compartment for products of these genes (4.8×10^−76^). The enrichment of this set of high-*S*_het_ genes for transcription factors is consistent with literature that describes dosage dependence for enzymatic proteins and haploinsufficiency for transcriptional regulators^36^.

**Figure 6:**
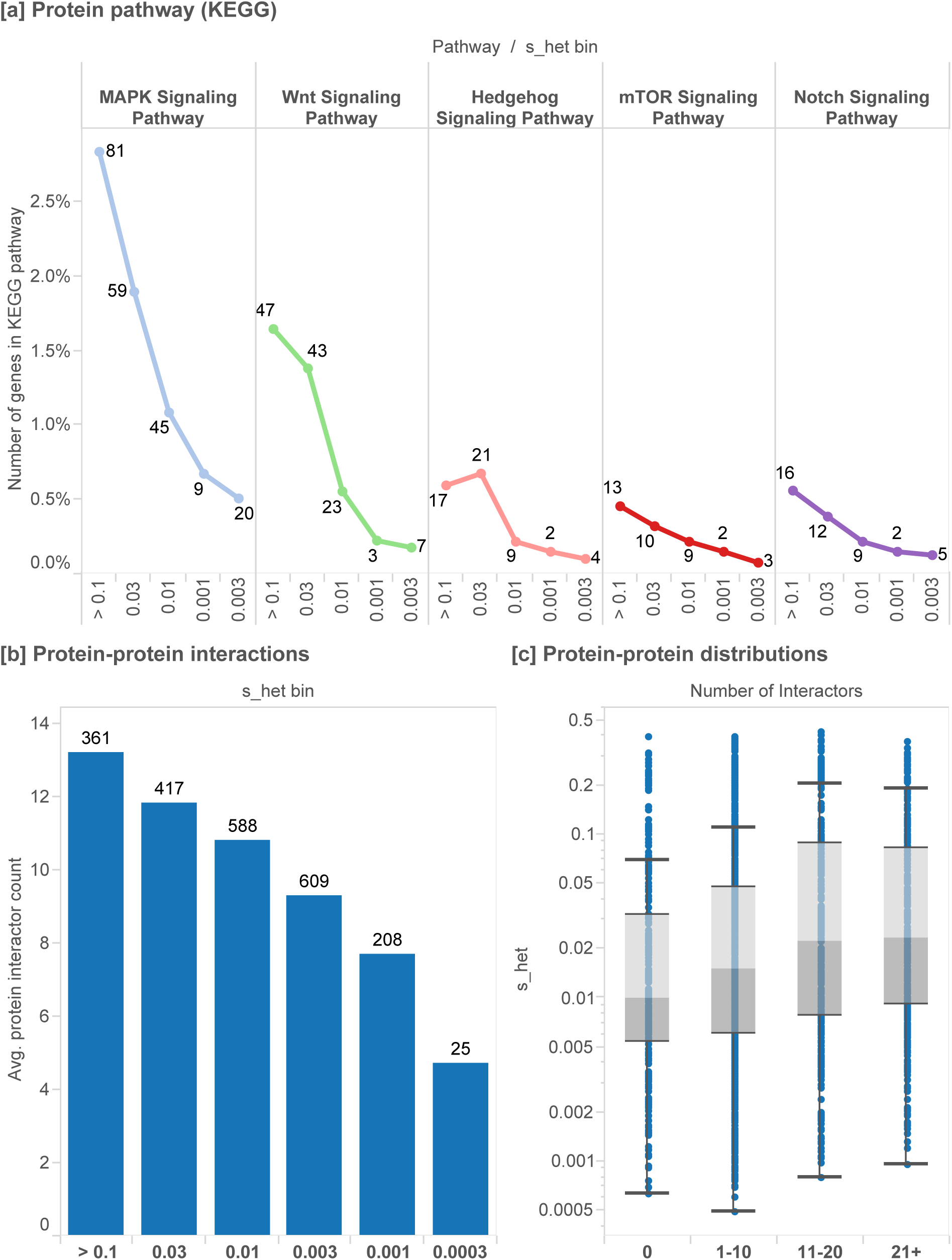
Protein pathways and protein-protein interactions. [a] In key developmental pathways in KEGG, we find that genes with higher *S*_het_ values are enriched in genes important to development. [b] We plot the distribution of the number of protein-protein interactions for each gene, as determined by a genome-wide Mass Spectrometry assay^37^ versus *S*_het_ value. [c] We find that *S*_het_ values are positively correlated with the number of observed interactors for each gene. Box plots range from 25^th^-75^th^ percentile values and whiskers include 1.5 times the interquartile range.

Estimation of the strength of purifying selection on PTVs provides a measure of gene dispensability unbiased with respect to existing knowledge. Thus, it has the potential to highlight genes that play a key role in development or in maintaining core functions in human cells. There are many genes with high estimated fitness costs that have not been previously described in human genetics studies. Given the marked enrichment of genes with high *S*_het_ values associated with Mendelian disorders, cell essentiality, embryonic lethality and development, it is plausible that many genes with high *S*_het_ values that have not been previously associated with human disease may be so detrimental that they are required for embryonic development.

We inspect the set of genes that lack disease annotations and publications but that have high *S*_het_ values to determine whether they share functional and genetic features reminiscent of known genes with central roles in cell housekeeping and developmental biology. We measure the relative knowledge about each gene in the primary literature from Entrez and PubMed^38^ using the number of gene reports connected with each manuscript, and sum the weighted contributions across all available manuscripts^39^ (PubMed score, **Methods**). While the PubMed score is positively correlated with *S*_het_ values, a substantial number of understudied genes fall in the highest *S*_het_ decile (**Supplementary Figure 6**).

We selected the 250 most cited and least cited genes within the top *S*_het_ decile, and compared their frequency of protein-protein interactions, viability of orthologous mouse knockouts and cell essentiality assays. Genes with the fewest publications have nearly the same number of embryonic lethal mouse knockouts as genes with the most publications. Other assays are only slightly depleted in the set of genes with the fewest publications (**Supplementary Figure 7**). These findings suggest that there may be additional essential developmental pathways yet to be uncovered in the set of genes under strong selection that lack functional or disease annotations, and provides a very promising gene set for further exploration. We have created a prioritized list of genes using a heuristic score developed from functional evidence to indicate the most promising candidates for future functional screening (**Supplementary Table 4**).

To place our inferences in the broader evolutionary context, we use comparable estimates from model organisms including flies and yeast, based on knockout competition with wild type or explicit crosses. In yeast, the analysis of a library of PTV knockouts provides a mean estimate of *S*_het_ ≈ 0.013, which is close to our inferred results (*S*_het_ ≈ 0.059) in humans^40^, given that the functional experiments excluded genes with very high *s*. Estimates in flies derived from homozygote lethal mutations which reduce viability in heterozygotes (rather than only PTVs) suggest values of *S*_het_ on the order of 1-3%, which is also in broad agreement with our estimates in humans^1,41^. While values of *s* in this range have a small impact in each generation, they may have dramatic evolutionary consequences^42^.

In conclusion, we use the genome-wide distribution of PTVs to estimate the fitness loss due to the heterozygous loss of each gene. Unlike recent work on intolerance to variation and its utility in human genetics^19,43^, we attempt to explicitly estimate the distribution of selection coefficients for PTVs. Our estimates are also distinct from the earlier work on the estimation of fitness effects of allelic variants in humans^44^ as the large sample size coupled with the assumption of strong selection makes our approach robust with respect to complexities of demographic history and dominance, and allows gene-based inferences. Conversely, our assumptions are justified for many but not all genes, as the method has limited resolution for genes under the strongest and weakest selection. We find significant enrichments in genes under strong selection in orthologous lethal mouse knockouts, genes that are essential for cell proliferation, and transcription factors. Additionally, these results may be useful in Mendelian disease gene discovery efforts and provide clinical utility in the inference of severity and mode of inheritance underlying Mendelian disease.

## Methods

### Model of deterministic mutation-selection balance

For most genes, protein-truncating alleles are both individually and collectively rare. For genes where they are collectively rare, estimation of the selective effect against heterozygous PTVs (*S*_het_) can be greatly simplified. We model each gene as a single bi-allelic locus with cumulative frequency *X* = ∑ _*j*_*x*_*j*_ where the sum is over PTVs in gene *i* for PTV sites *j*. This is motivated by the simplifying assumption of identical selection coefficients for all PTVs within a gene, and the observation that the frequency of the vast majority of PTVs is extremely low such that the occurrence of multiple variable sites within a gene on a single haplotype is also extremely low (2*Nx*_*ij*_*x*_*ik*_ < 1 for sample size *N*). Moreover, multiple PTVs in a gene in an individual would be functionally equivalent to a single PTV resulting in a loss of function state.

Then for each gene, the cumulative allele frequency *X* is influenced by incoming mutation, selection and the random reassortment of alleles (genetic drift). When selection is strong, *S* ≫ 2.5×10^−5^ (i.e. when 4*N*_e_*S* ≫ 1, with effective population size *N*_e_ = 10^4^), drift is much smaller than the contribution of selection. Furthermore, the strength of genetic drift is weakest for genes at low frequencies: for a variant with cumulative frequency of *X* = 0.001 the expected frequency change due to drift is only 〈Δ*X*^2^〉 ~ *X*/4*N*_e_ = 2.5×10^−8^ per generation. Notably, at the locus level assuming *X* ≪ 1 the drift contribution is also much smaller than the mutational influx. Hence under strong selection and for small allele frequencies the expected cumulative frequency of PTVs is determined by the equilibrium between the influx of *de novo* mutations (estimated to increase the cumulative frequency by an average 1.4×10^−6^ per locus per generation by mutational model) and the outflux due to natural selection.

In the presence of selection on both heterozygotes and homozygotes and ignoring back mutations, the dynamics of *X* are captured by the following equation:

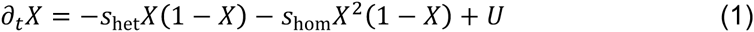

Here *U* represents the PTV mutation rate at the gene locus per individual per generation, and *S*_het_ = *hS* > 0 and *S*_hom_ = *S* > 0 represent the strength of negative selection against PTV heterozygotes and homozygotes, respectively. We note that compound heterozygotes (with a single PTV on each chromosome) are treated as homozygotes under the bi-allelic assumption. Provided *X* ≪ 1, as is the case for PTVs under strong selection (2*N*_e_*S* ≫ 1), this equation simplifies dramatically:

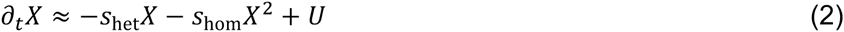

Because *X* ≪ 1, selection against heterozygotes (the linear term) generally also dominates over selection against homozygotes (the quadratic term), provided *S*_het_/*S*_hom_ ≫ *X*. This is only violated in cases of extreme recessivity (where the dominance coefficient *h* ≪ 0.001), but even in that case the expected cumulative frequency of PTVs in essential genes is unlikely to exceed 0.001 (the characteristic *X* in the completely recessive case is 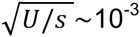 when *S*~1, see simulations in **Supplementary Figure 1**). The strong selection regime thus corresponds to mutation-selection balance in the heterozygote state of a PTV mutation. Notably, there is no dependence on the demography or population size in this regime, as the contribution from drift vanishes because selection drives alleles out of the population efficiently and on very short time scales.

From Eq. 2 follows that for a population sample of size *N*, sample allele counts 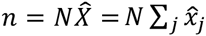 are expected to be Poisson distributed around the expectation given by:

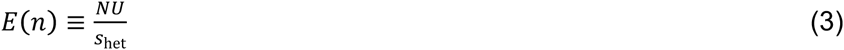

Generally, genes under the strongest and weakest selection are expected to have greater estimation uncertainty, as the resolution to estimate *S*_het_ deteriorates when variants are so common that they may not only be controlled by heterozygote selection, but also by drift or complex demography. However, the overwhelming majority of genes conform to our assumptions of cumulative PTV allele frequency not exceeding 0.001. Despite issues such as the admixture of populations, consanguineous samples in ExAC^24^, and the Wahlund effect, very few genes (1,201 of 17,199 covered genes) have higher estimated cumulative allele frequencies 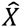, which we restrict from the estimation procedure. On the other end of the spectrum, genes under strong selection may lack PTVs by chance alone in ExAC, which limits the ability to distinguish between large selective effects.

### Population genetics simulations of model assumptions

To validate the assumption that estimates of selection can be made under mutation-selection balance independent of demography or population size for variants under sufficiently strong selection, we used SLiM 2.0 to conduct forward population genetics simulations^45^. We ran 10 replicates each of simulations with selection coefficients of −5×10^−2^, −5×10^−3^, −5×10^−4^, −5×10^−5^, and −5×10^−6^ through the demography published in Tennessen *et al.*^46^ for Africans and Europeans (**Supplementary Figure 1**). We compare the theoretical mutation load (*U*/*S*_het_) with the simulated mutation load in three groups (African, European and Combined, which includes pooled site frequency spectra from both African and European populations in proportions represented in the EXAC dataset). The simulations support our assumption of mutation-selection balance in the strong selection regime (|*S*_het_| >= 1×10^−3^), which appears to be appropriate for PTVs.

All simulations had a length of 1 kilobase, mutation rate of 2×10^−8^ per generation per base pair, and recombination rate of 1×10^−5^ per generation per base pair. The high recombination rate was chosen to simulate largely unlinked sites, as we are simulating PTVs which are infrequent enough that they are expected not to be in linkage with other PTVs in the same gene.

### Dataset for *S*_het_ estimation

In this analysis, we use Exome Aggregation Consortium (ExAC) dataset version 0.3, a set of jointly-called exomes from 60,706 individuals ascertained with no known severe, early-onset Mendelian disorders. The mean coverage depth was calculated for each gene (canonical transcript from Ensembl v75, GENCODE v19) in the ExAC dataset (mean 57.75; s.d. 20.96).

Genes with average coverage depth of at least 30x were used in further analysis (N=17,199). Single nucleotide substitution variants annotated as PASS quality with predicted functional effects in the canonical transcript of stop-gain, splice donor, or splice acceptor (Variant Effect Predictor) were included in the analysis.

We are mindful that not all PTVs will result in complete loss of gene function, due to alternative transcripts or nonsense mediated decay. To address this, variants were filtered using LOFTEE^47^ and restricted to those predicted with high confidence to have consequences in the canonical transcript.

For each of the 17,199 genes we have observable values for (*n*, *U*, *N*), where *n* denotes the total number of observed PTV alleles in the population sample of *N* chromosomes covered in the gene, and *U* the PTV mutation rate across the canonical gene transcript from a mutational model^19,20^. Values of *N* and *U* for each gene from Samocha *et. al.* were used along with the number of well-covered chromosomes in each gene to generate the null mutational expectation of neutral evolution, *NU*. Incorrectly specified values from this mutational model could alter estimates of selection for individual genes, as higher estimates of selection are made in genes with greater depletions from the null expectation model. Our inference of selection coefficients relies on the assumption that the cumulative population frequency of PTV mutations, *X*, is small due to strong negative selection, so genes with 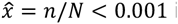 are omitted from the analysis, leaving 15,998 genes.

### Estimation of *P*(*S*_het_)

A genome-wide ensemble of observed (*n*) and expected (*NU* ≡ *v*) genic PTV counts enables the inference of the distribution of heterozygous loss-of-function fitness effects, *P*(*S*_het_), which underlies the evolutionary dynamics of this class of mutations. We estimate the parameters (α, β) of this distribution by fitting the observed distribution of PTV counts across genes:

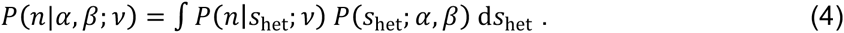

For a given gene under negative selection PTV mutations are rare events, such that we expect a Poisson distribution for the likelihood of the observed number of PTVs

*P*(*n*|*S*_het_;*v*) = Poiss *n*; λ), where λ = *v*/*S*_het_ (Eq. 3). We parameterize by using the functional form of an inverse Gaussian distribution, i.e. *P* (*S*_het_; α, β = IG(*S*_het_; α, β), so Eq. 4 becomes:

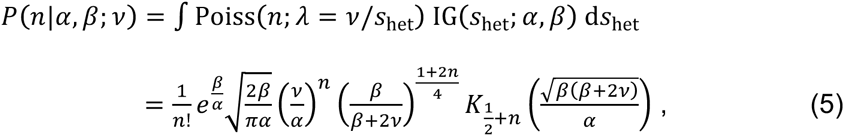

where *K*_*n*_(*Z*) is the modified Bessel function of the second kind. To estimate parameters of the distribution of selection coefficients, *P*(*S*_het_; α, β), we fit Eq. 5 to the observed distribution of PTV counts, *Qn*, by maximizing the log-likelihood

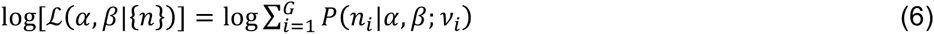

on the regime α ∈ [10^−3^, 2] and β ∈ [10^−4^, 2], where *G* is the number of genes. In order to account for a slight positive correlation between the mutation rate and selection strength (**Supplementary Figure 8**), we separately perform the fit on *U* terciles of the data set and combine the results in a mixture distribution with equal weights. The mean mutation rates in the three terciles are 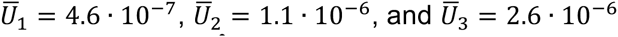. We estimate 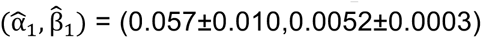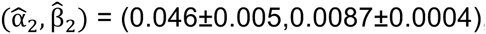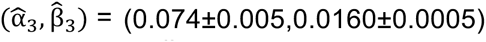, with error margins denoting two s.d. from 100 bootstrapping replicates of the set of ~5,333 genes in each tercile. This error estimate is intended to quantify the effect of the sampling noise in the data set on the parameter inference while local mutation rate estimates are assumed fixed. The resulting fitted distributions of counts are shown in **Supplementary Figure 9** together with the corresponding *Q* (*n*), while **Figure 1** shows the inferred 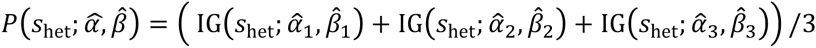. The choice for the functional form of *P* (*S*_het_) is motivated by the shape of the empirical distribution of the naïve estimator *v*/*n* (given by a simple inversion of Eq. 3). We also compared the log-likelihood of the fit to *Q*(*n*) obtained with this model to that obtained from two other two-parameter distributions, *S*_het_ ~ Gamma and *S*_het_ ~ InvGamma, and chose the model with the highest likelihood, which is *S*_het_ ~ IG.

### Inference of *S*_het_ on individual genes

From the inferred distributions 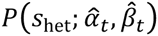 in each tercile *t* of the mutation rate *U*, we construct a per-gene estimator of *S*_het_ for genes in the tercile using the posterior probability given *n*, which mitigates the stochasticity of the observed PTV count:

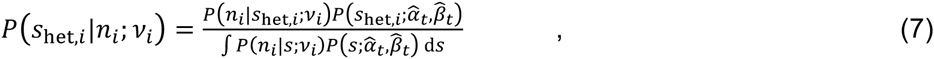

where the denominator is given by Eq. 5. **Supplementary Table 1** provides the mean values derived from these posterior probabilities for each gene.

### Predicted mode of inheritance in clinical exome cases

We trained a Naïve Bayes classifier to predict the mode of inheritance in a set of solved clinical exome sequencing cases from Baylor College of Medicine (N=283 cases)^22^ and UCLA^23^ (N=176 cases). Using data from UCLA as the training dataset, we are able to cross-predict the mode of inheritance in separately ascertained Baylor cases with classification accuracy of 88.0%, sensitivity of 86.1%, specificity of 90.2%, and an AUC of 0.931. Genes that were related to diagnosis in both clinics (overlapping genes) were removed from the larger Baylor set (**Supplementary Figure 2**).

Using a logistic regression based on the full set of cases from Baylor and UCLA, we generated predictions for all 15,998 genes where there is a *S*_het_ value (**Supplementary Table 4**).

### Mouse knockout comparative analysis

We reviewed mouse knockout enrichments from two datasets: the full set of mouse knockouts from a neutrally-ascertained mouse knockout screen (N=2,179 genes) generated by the International Mouse Phenotyping Consortium^25^. Genes were classified as ‘Viable’, ‘Sub-Viable’, or ‘Lethal’ based on the results for the assay.

### PubMed gene score and enrichment analysis

We developed a score to estimate the relative importance of each gene in the published medical and scientific literature. First, we connected literature from Entrez which included both PubMed citations and references to Entrez genes. We assigned a weight to each article referencing a gene of 1/*a_i_*, where *a_i_* was the number of genes referred to by article *i*. For example, an article referring to four genes would receive a weight of 1/4. Finally, we assigned each gene a score which was the sum of the weighted article scores. These scores ranged from 4,672 articles per gene (p53) to 0.0001 articles/gene.

Next, we focused on genes that are estimated to be under very strong selection but that lack functional or clinical annotations. In the top decile of *S*_het_ values, we separated the top 250 and bottom 250 genes by PubMed score. We then annotated each of these with unbiased genome-wide assays, including the number of protein-protein interactions (as determined by a genome-wide Mass Spectrometry assay)^37^, whether each gene is determined to be cell-essential in genome-wide CRISPR and gene trap assays^3^, and whether there is a mouse knockout in the neutrally-ascertained orthologous nonviable mouse knockout^48^. To limit the number of genes with incorrect *S*_het_ estimates in this set of 500 genes, we pre-filtered any genes with only a single exon, as they may be enriched for recent pseudogenes, and also removed any olfactory, mucin, and zinc finger proteins.

### Functional enrichment analysis

We inspected the functional annotations related to approximately the top 10% of selectively disadvantageous genes (with *S*_het_ > 0.15, N=2,072 genes) that were successfully mapped using Database for Annotation, Visualization, and Integrated Discovery (DAVID) version 6.7^35^, DAVID. Separately, two other cutoffs (*S*_het_ > 0.25, N=897 genes and *S*_het_ > 0.5, N=32 genes) were also tested and similar results were identified.

Using DAVID, we identified functional annotation terms and keywords that were enriched and clustered. Functional annotation terms were generated using the Functional Annotation tool, which includes protein information resource keywords, GeneOntology (GO) terms, biological processes and pathways, and protein domains. Using the default settings (Count 2 and EASE 0.1), 247 statistically significant (Bonferroni corrected) terms were identified and are included in **Supplementary Table 5.**

Using the DAVID Functional Annotation clustering feature, we identified clusters using the same set of 2,072 genes with the default settings. The first annotation cluster includes core, essential cellular components including the nuclear lumen, nucleoplasm, organelle lumen (Enrichment score 32.63), and the second includes transcription regulation and transcription factor activity (Enrichment score 27.94), detailed in **Supplementary Table 6.**

## Acknowledgements

This work was supported by National Institutes of Health Grant HG007229, GM078598, HG009088, MH101244, and GM105857. We thank Ivan Adzhubei, Konrad Karczewski, and Alexey Kondrashov for helpful advice.

## Supplementary Information

**Supplementary Figure 1:**
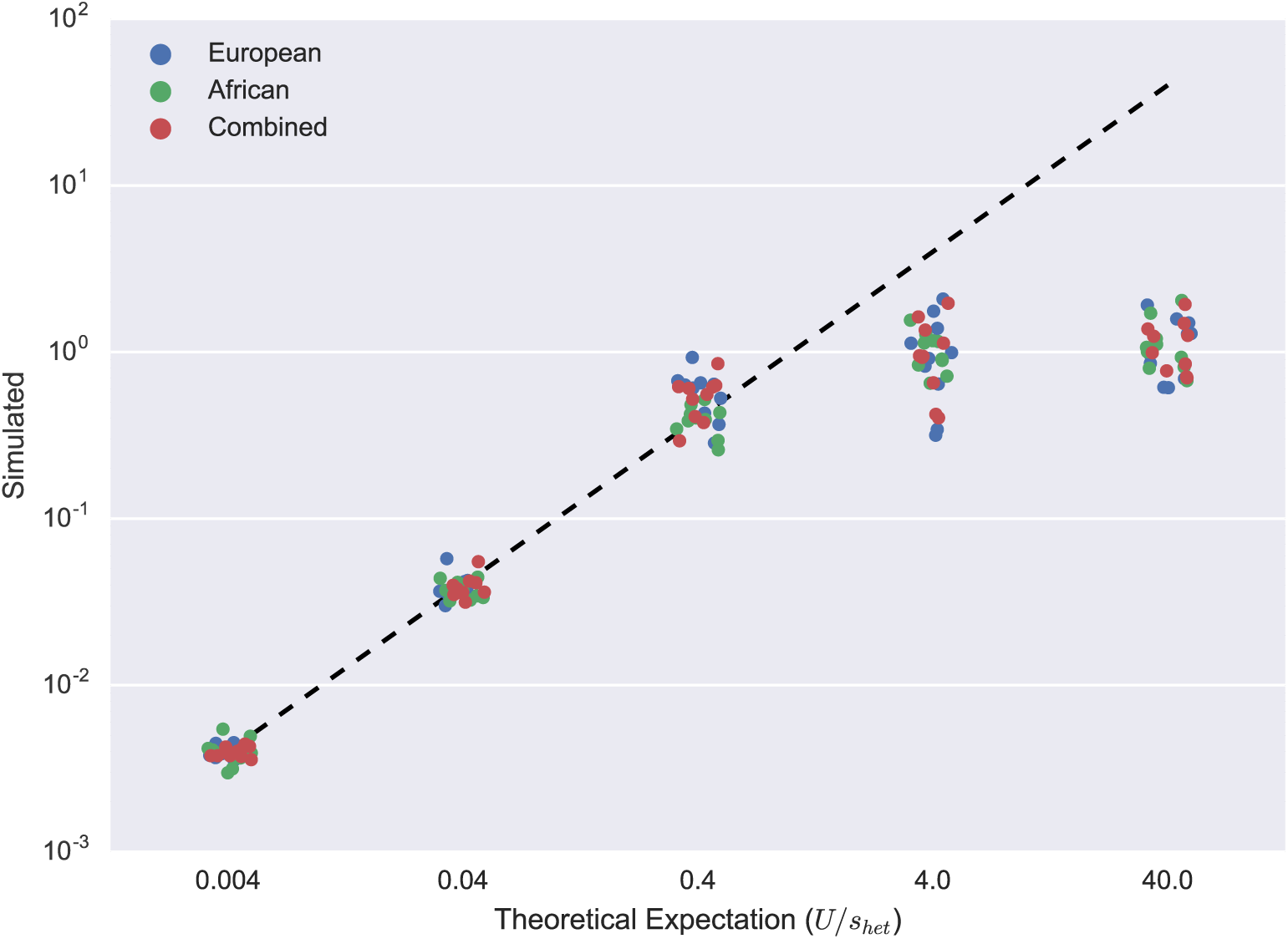
**Population genetics simulations of model assumptions.** To validate the assumption that estimates of selection can be made under mutation-selection balance independent of demography or population size for variants under sufficiently strong selection (**Methods**), we used SLiM 2.0 to conduct forward population genetics simulations^45^. We compare the theoretical mutation load (*U*/*S*_het_) with the simulated mutation load in three groups (African, European and Combined, which includes pooled site frequency spectra from both African and European populations in proportions represented in the EXAC dataset) for *S*_het_ ∈ {−5×10^−2^, −5×10^−3^, −5×10^−4^, −5×10^−5^, −5×10^−6^} from left to right on the x-axis. *U* = 2×10^−3^ for all simulations. The simulations support our assumption of mutation-selection balance in the strong selection regime (|*S*_het_| > 1×10^−3^), which appears to be appropriate for PTVs. A small random jitter has been added to separate the points visually.

**Supplementary Figure 2:**
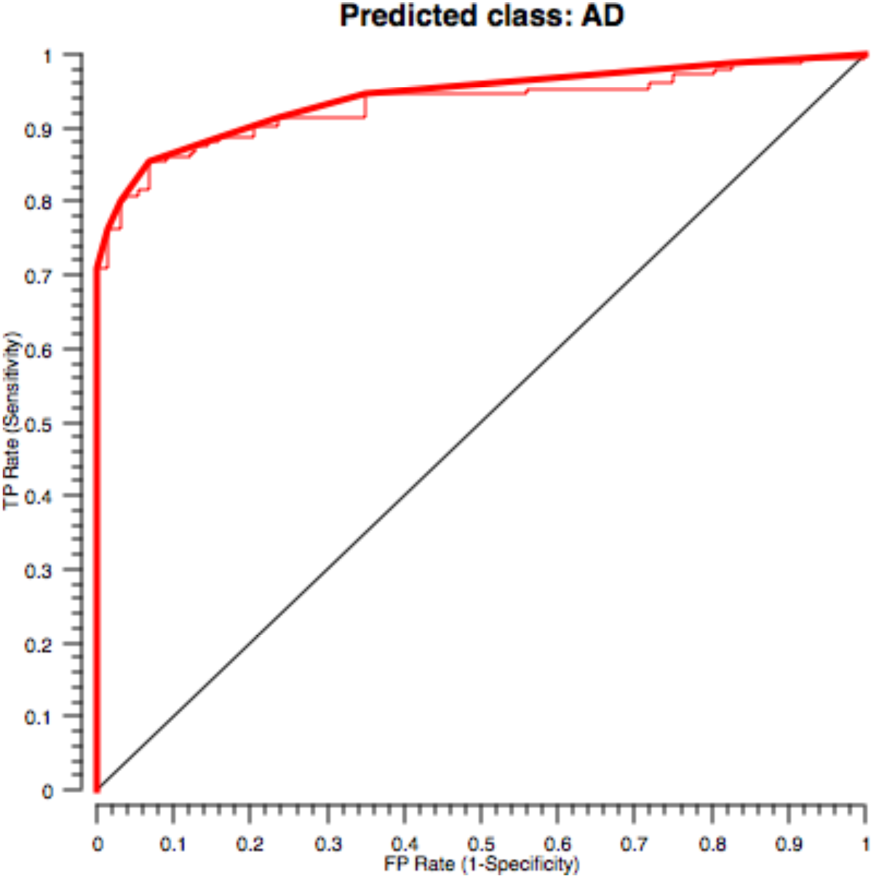
**Mode of inheritance gene classifier performance characteristics ROC curve for mode of inheritance gene classifier.** We train a Naïve Bayes classifier to predict the mode of inheritance in a set of solved clinical exome sequencing cases from Baylor College of Medicine^22^ (N=283 cases) and UCLA^23^ (N=176 cases). Using data from UCLA as the training dataset, we are able to cross-predict the mode of inheritance in separately ascertained Baylor cases with classification accuracy of 88.0%, sensitivity of 86.1%, specificity of 90.2%, and an AUC of 0.931. Genes that were related to diagnosis in both clinics (overlapping genes) were removed from the larger Baylor set.

**Supplementary Figure 3:**
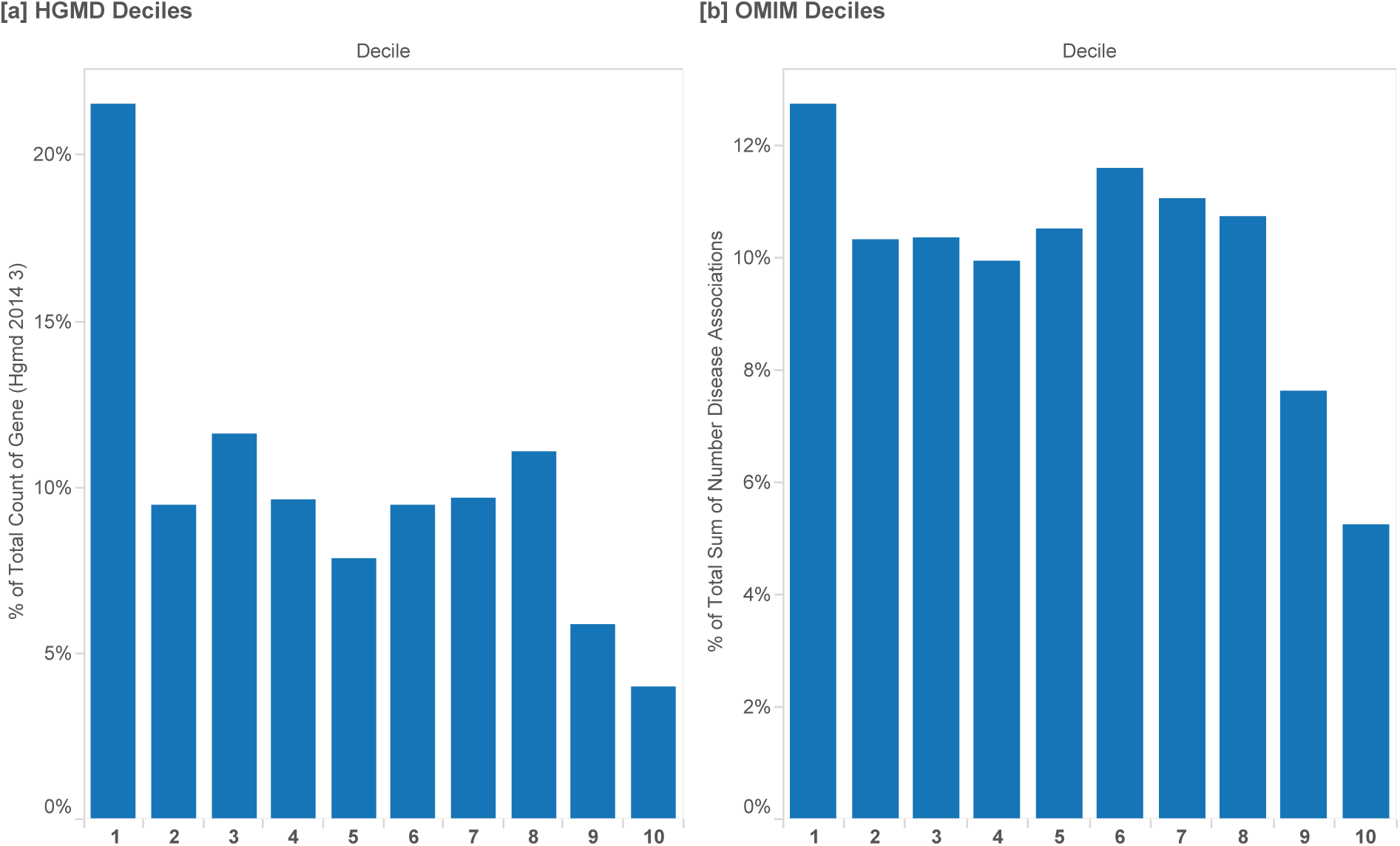
**HGMD and OMIM disease annotations** We next annotate each gene with disease associations using reports in the medical and scientific literature in the Human Gene Mutation Database (HGMD v.2014_3) and also with the number of associations listed in the Online Mendelian Inheritance in Man (OMIM MorbidMap)^49^. HGMD disease gene annotations (DM status) are used to specify a suspected association with a known Mendelian disorder, and all associations from OMIM are included, although it is well-known that these disease annotations have quality and curation issues^50,51^. Data are separated by *S*_het_ deciles and OMIM and HGMD disease annotations appear to be a combination of Autosomal Dominant and Autosomal Recessive disease genes.

**Supplementary Figure 4:**
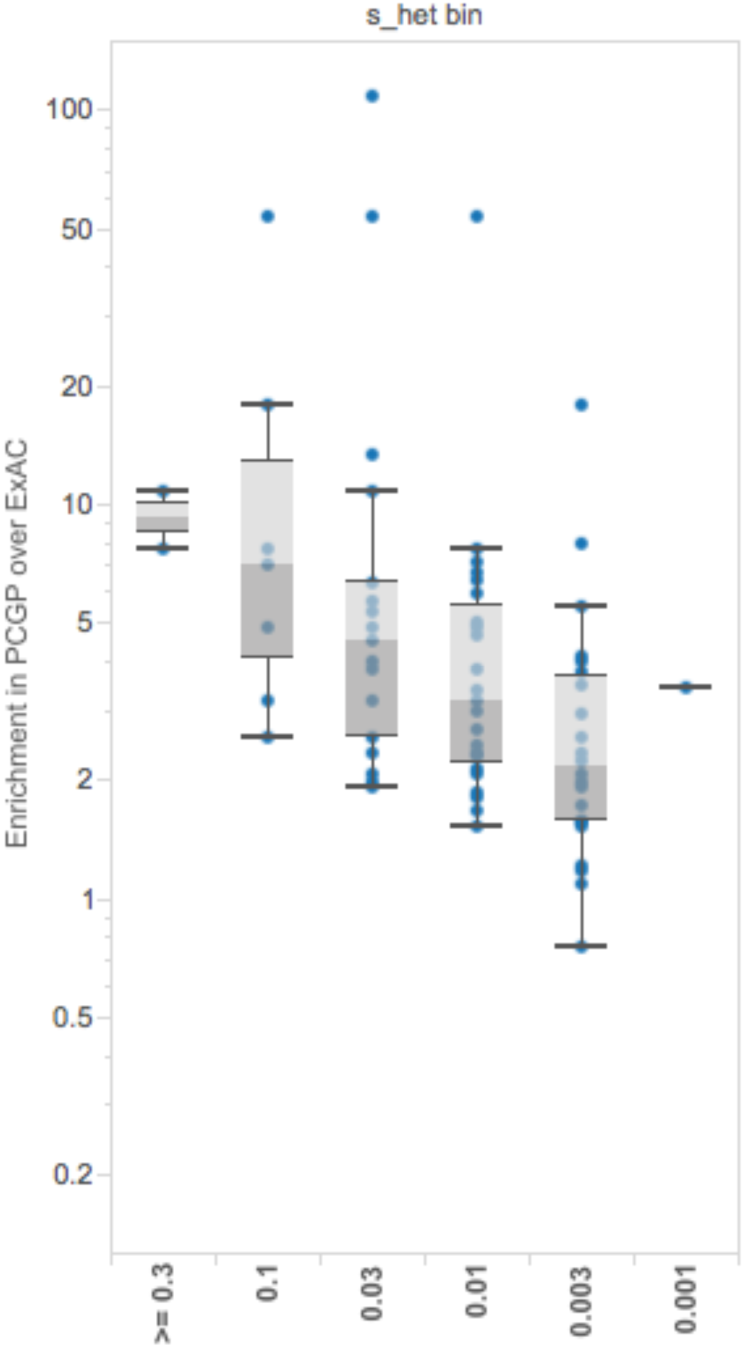
**Enrichment in germline cancer predisposition genes Enrichment in germline cancer predisposition genes.** In a large screen of germline cancer predisposition genes in the Pediatric Cancer Genome Project (PCGP), the enrichment of variants in pediatric cancer cases is measured over individuals in ExAC.^52^ Genes with greater enrichment of variants in cancer cases over ExAC are correlated with higher selection coefficients. Data are separated by *S*_het_ bins on a log scale. Box plots range from 25^th^–75^th^ percentile values and whiskers include 1.5 times the interquartile range.

**Supplementary Figure 5:**
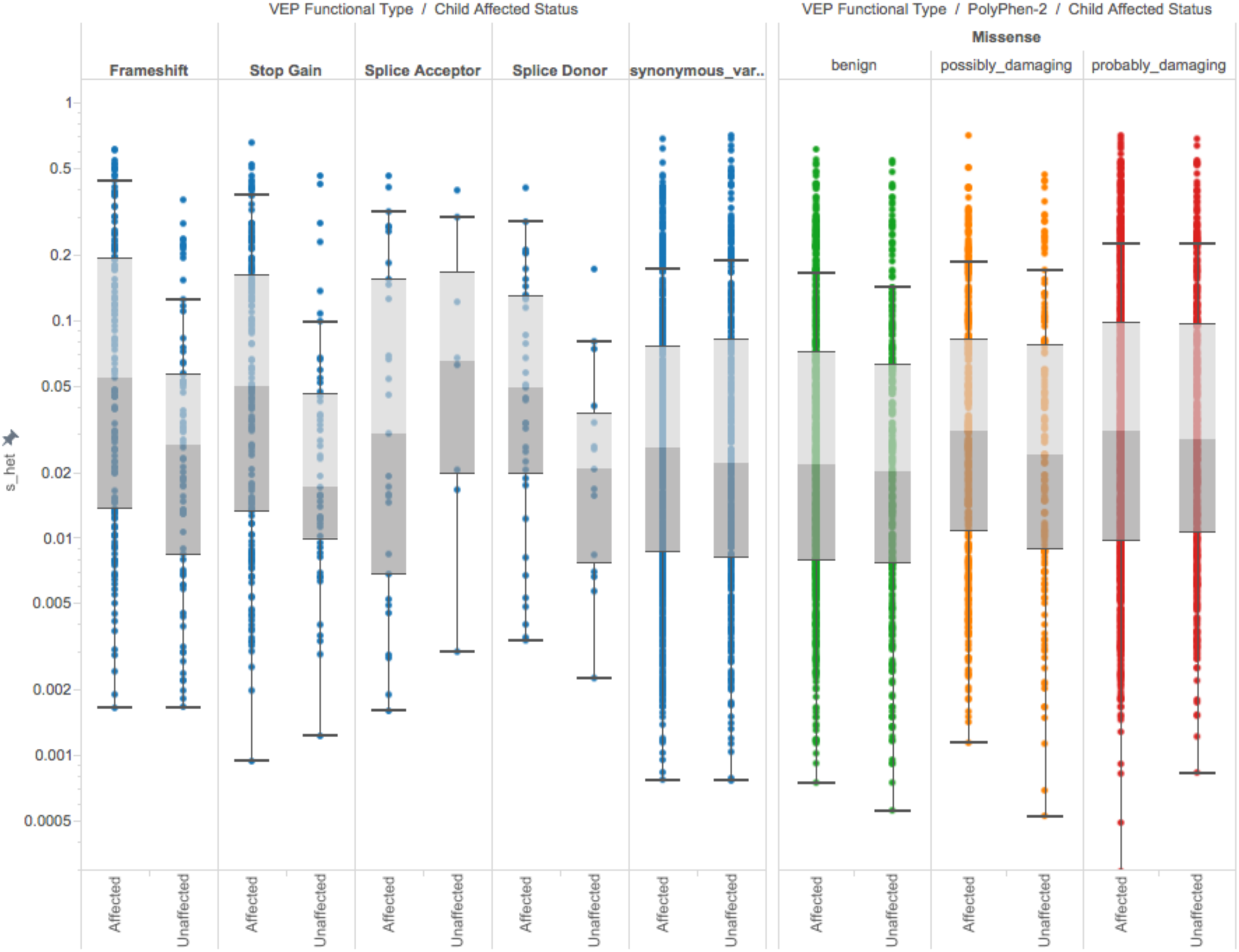
**Enrichments of** *S*_het_ **in *de novo* variants in autism spectrum disorders (ASD) Enrichments of** *S*_het_ **in *de novo* variants from autism spectrum disorders (ASD) case and control trios.** In a set of *de novo* ASD case (N=2,939) and control (N=1,429) trios, *S*_het_ estimates can help discriminate between all protein-coding variants, protein-truncating variants (including all frameshift, nonsense, and essential splice site variants), and individually for nonsense, frameshift, and missense variants which are predicted to be PolyPhen-2 damaging. Box plots range from 25^th^-75^th^ percentile values and whiskers include 1.5 times the interquartile range.

**Supplementary Figure 6:**
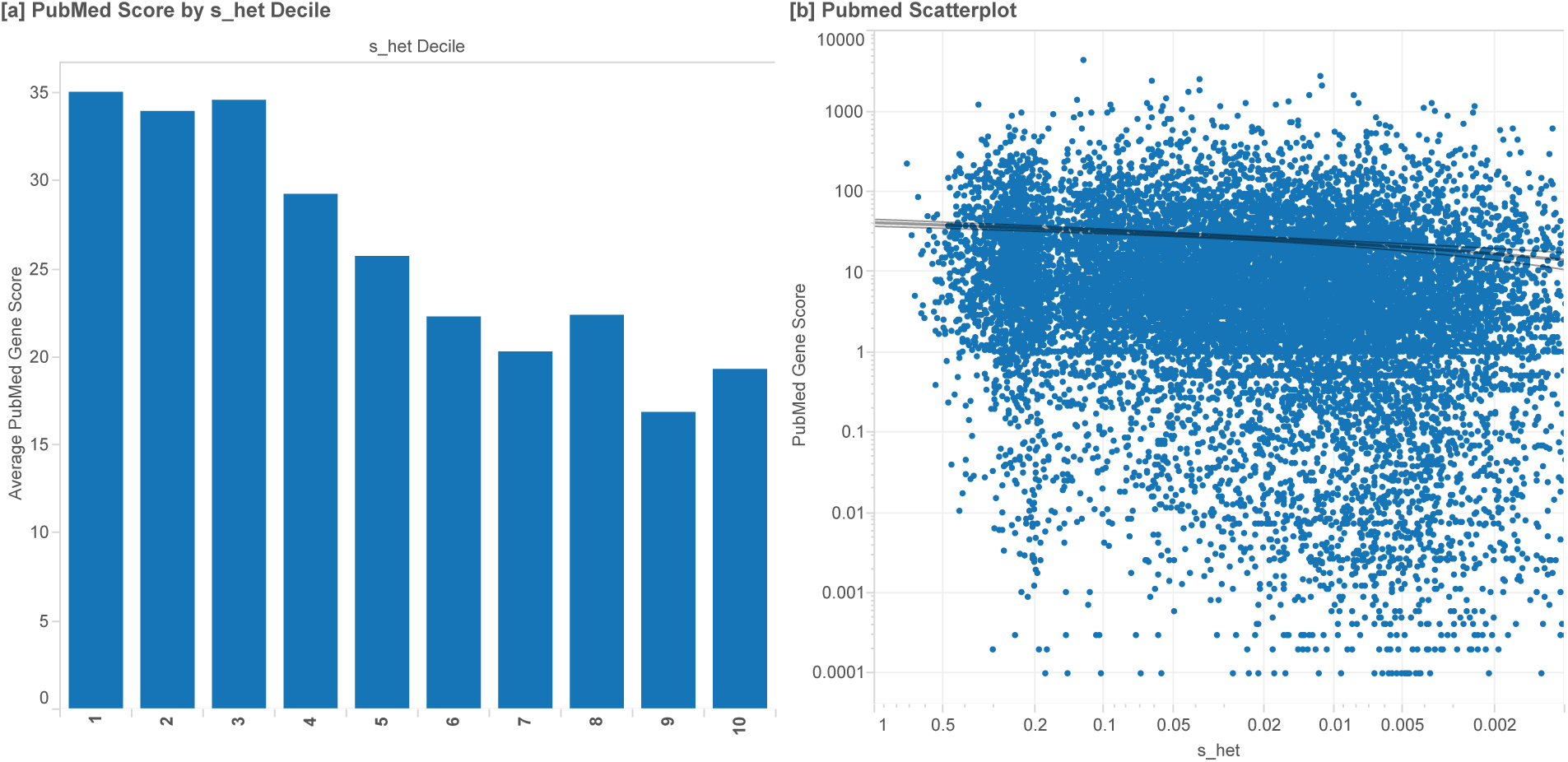
**PubMed score** We measured the relative knowledge about each gene in the primary literature using Entrez and PubMed^38^. We developed a novel PubMed Gene Score using the number of reports connected with each gene and the number of genes described in each manuscript, and sum the relative contributions for each gene across all available manuscripts. The Pubmed Gene Score is significantly positively correlated with *S*_het_ (p<0.0001), so we know that the average knowledge has a positive relationship with gene selection/consequence of disruption. We also find that there are a substantial number of genes with high predictions of *S*_het_ but not even one full, individual citation in the scientific literature. **Association of** *S*_het_ **estimates with PubMed gene score.** [a] The average PubMed gene score is calculated by *S*_het_ decile. Estimates of selection are positively correlated with the average Pubmed gene score. Each bin contains 10% of all covered genes, ordered from greatest to smallest *S*_het_ values, in bins 1 through 10, respectively. [b] The PubMed gene score is significantly positively correlated with the (p<0.0001) using a logarithmic model (y=4.557*log(*S*_het_)+44.449) with R^2^=0.00409.

**Supplementary Figure 7:**
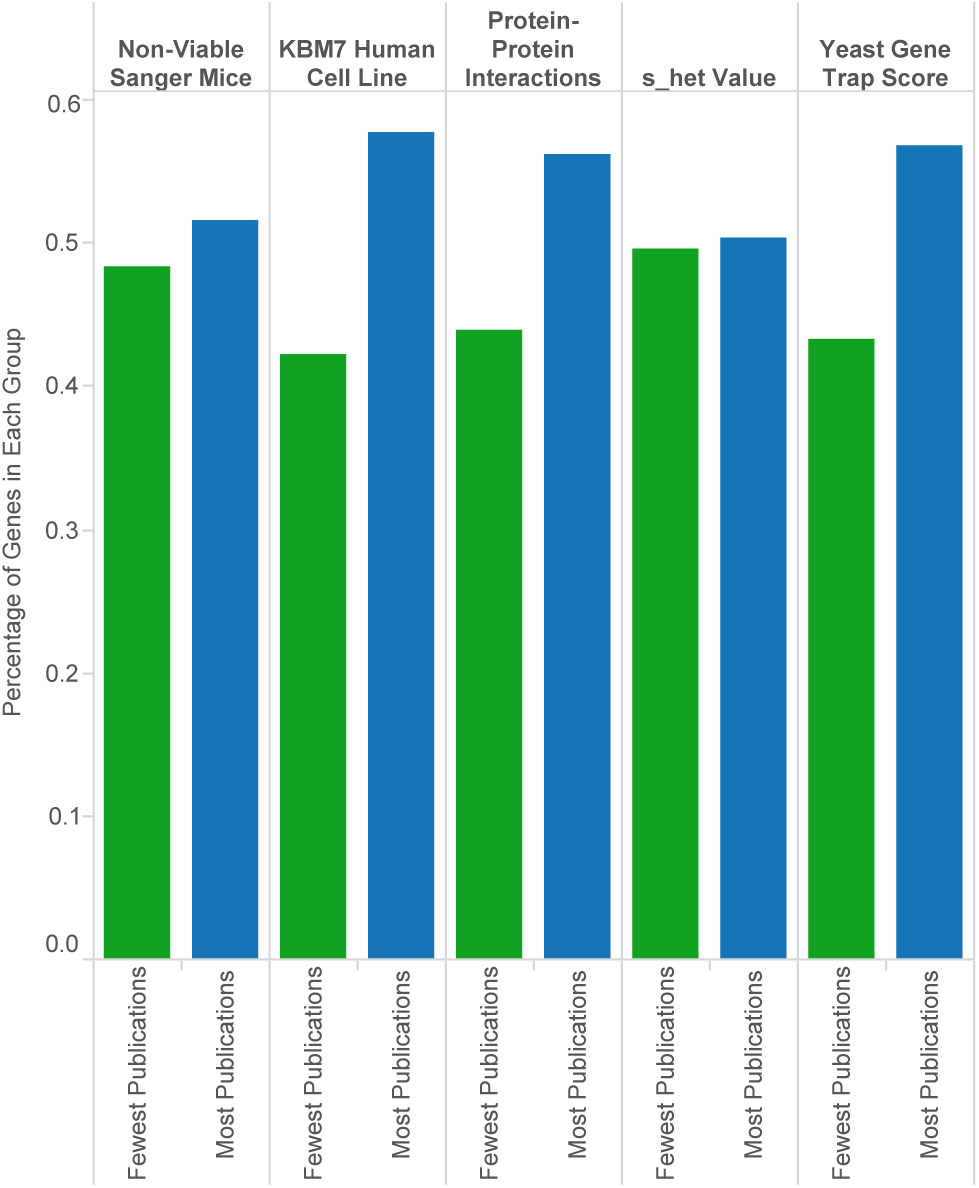
**Most published and least published genes from top** *S*_het_ **decile Most published and least published genes from top** *S*_het_ **decile.** The proportion of annotations related to genes with the fewest and most publications in Entrez Gene. From the set of genes under the strongest selection (top 10% of *S*_het_ values), we create two sets of 250 genes. The first set of genes has the fewest publications associated with each gene, as defined by our PubMed gene score (**Methods**), and the second set has the greatest number of associated publications. Between the two groups, we compare the *S*_het_ values, number of protein-protein interactions, viability of orthologous mouse knockouts (IMPC), and cell essentiality assays (KBM-7 CRISPR score and Gene Trap Score). These results suggest that the genes in the least published set are similar to those in the most published set, and are also potentially important developmental genes.

**Supplementary Figure 8:**
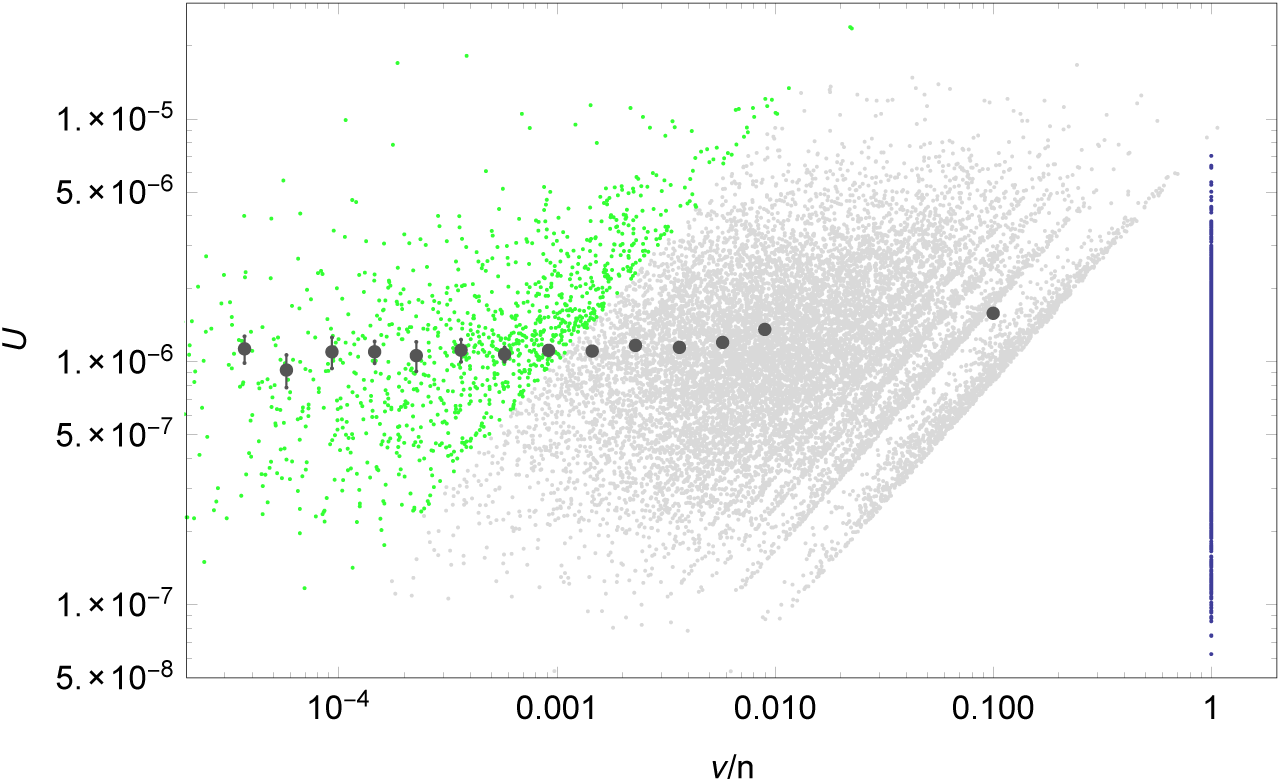
**Relationship between gene mutation rate and selection Relationship between gene mutation rate and selection.** Relationship between the estimate of local mutation rate, *U*, and the naïve estimator for heterozygous selection against PTVs,)*v*/*n* = *NU*/*n*, for all 17,199 genes. Light green dots represent genes with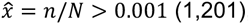, which we omit in the inference of the distribution of *P* (*S*_het_). Light gray dots are used genes with *n*>0 (14,274), while dark blue dots correspond to those with *n*=0 (1,724). The latter were assigned a fixed selection coefficient estimate of 1 for illustration purposes. We computed the mean *U* in logarithmic bins of)*v*/*n* for the range 0.00003 <*v*/*n* ≤ 0.012, and for the last bin from all genes with*v*/*n* > 0.012, including those with *n*=0 (large gray dots). Error bars denote s.e.m. The slight positive correlation between *U* and selection strength motivates the division of the data set into terciles of *U* and separate estimation of the parameters of the distribution of selection coefficients in each.

**Supplementary Figure 9:**
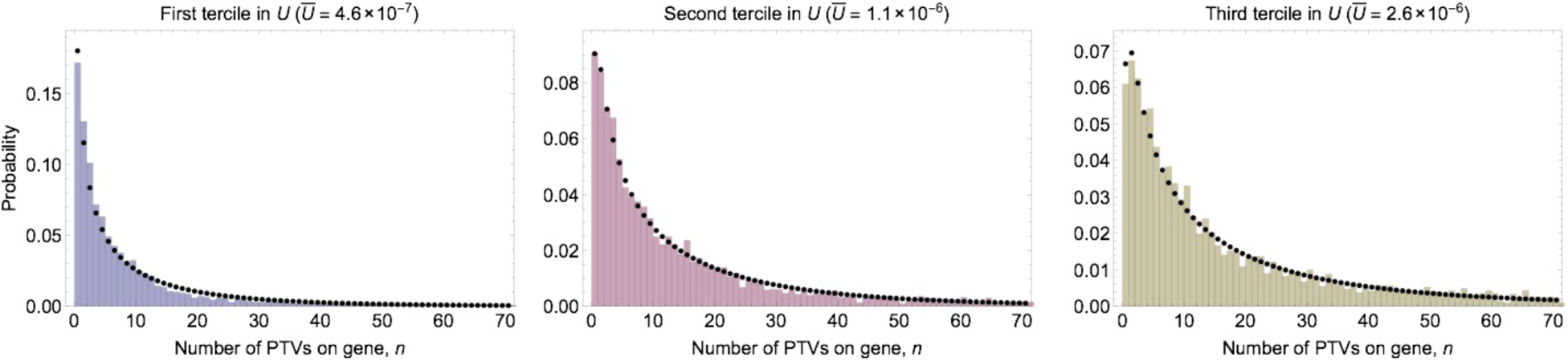
**Fit to the observed distribution of PTV counts Fit to the observed distribution of PTV counts.** Fitted distribution *P*(*n*) (black dots) from maximum likelihood fit to the observed distribution *Q*(*n*) (histogram) of PTV counts *n* across 15,998 considered genes divided into terciles according to mutation rate *U*, assuming 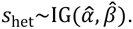.

**Supplementary Figure 10:**
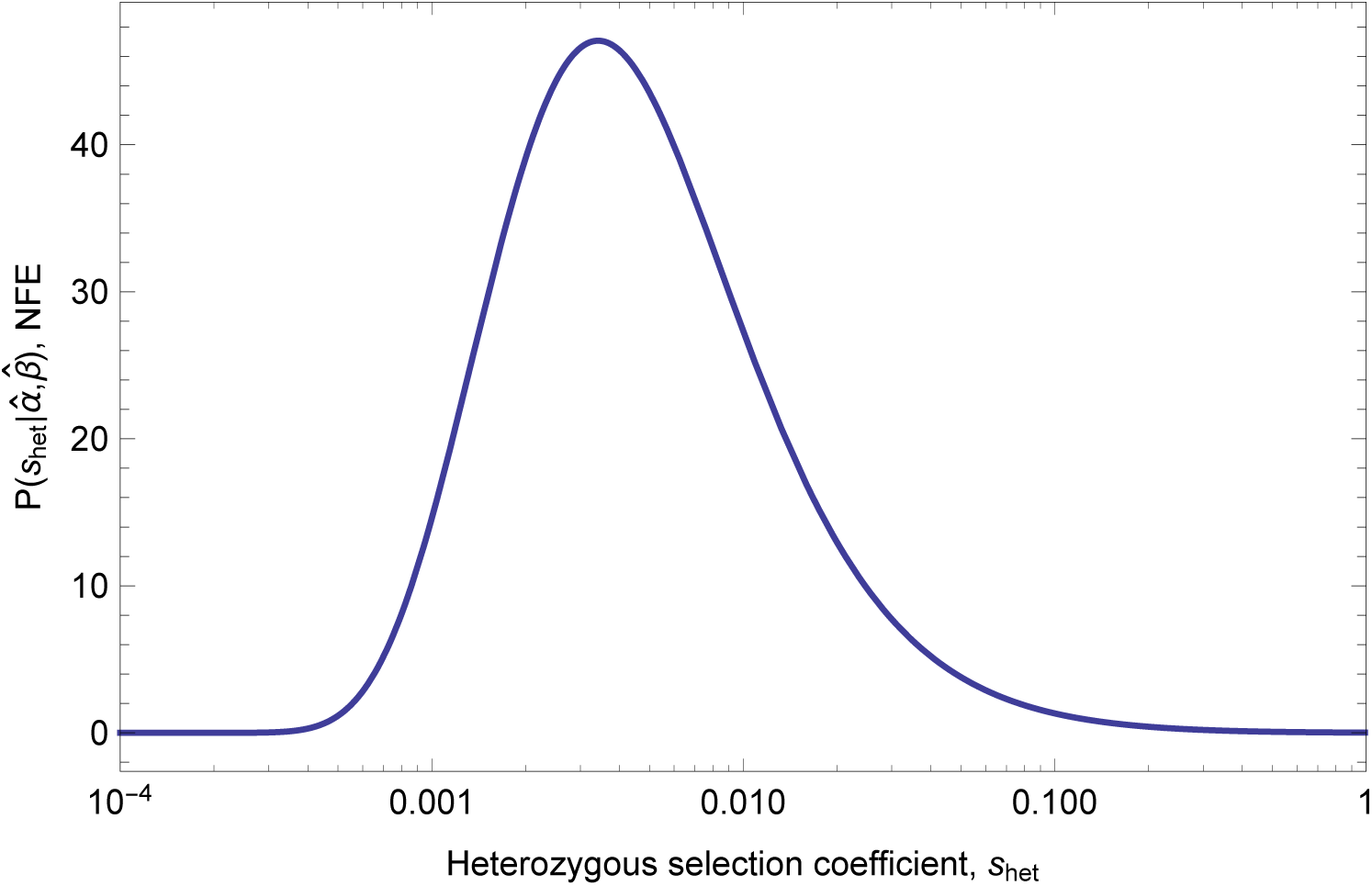
*S*_het_ **for non-Finnish Europeans** We separately repeated the inference procedure for *P* (*S*_het_) using data from a single population group, Non-Finnish Europeans (NFE, N=33,370, as annotated by ExAC), and generated a corresponding set of *S*_het_ estimates. The inferred parameters are very similar to those from the larger sample. Underlying allele count data is highly correlated between the overall set and the NFE set (R^2^ = 0.814) and the *S*_het_ estimates are also highly correlated. **Inferred distribution of fitness effects for heterozygous loss of gene function in non-Finnish Europeans.** Estimates of parameters from maximum likelihood fit to the observed distribution of PTV counts *n* across genes with 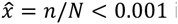 in the set of non-Finnish Europeans (16,279 genes), assuming *S*_het_~IG(α, β) in terciles of the mutation rate *U*. Parameter estimates are 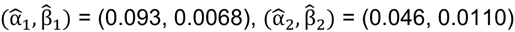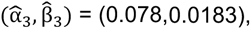, and shown is the mixture distribution of the three components with equal weights.

### Supplementary Note: Comparison with previous estimates of selection

Previous studies have assessed the functional importance of differential observed mutational load, and have shown that these aid in Mendelian and complex gene assessment. Here, we estimate the selective coefficient for heterozygous variants, rather than the overall depletion observed in each gene, using a Bayesian estimation approach. This approach mitigates power issues related to correlation with gene length, helps discriminate between dominant and recessive acting disease genes, and provides an estimate of an important evolutionary parameter (*S*_het_). These selective estimates can then be used to infer other related evolutionary parameters, such as dominance (*h*). Boyko *et al.*^5^ previously fit a gamma distribution to estimate the distribution of fitness effects for missense variants. Their results are grouped into three categories: *s <* 10^−4^ as neutral, 10^−4^ *< s <* 10^−2^ as moderately deleterious, and *s >* 10^−2^ as lethal, as there is insufficient resolution to distinguish between values above 0.01. The authors estimate that 30-40% of missense variants are in this lethal category. For PTVs, we find a larger number, 10,124 genes, or 63.2% of covered genes are estimated to have *S*_het_ > 0.01. In this study, we have limited resolution to accurately estimate *S*_het_ values between 0.15 and 0.34, and then poor resolution to differentiate specific *S*_het_ values above 0.34 (although they are estimated to be very high).

We also compared the *S*_het_ estimate with the ExAC pLI score and find that the overall correlation coefficient is *R^2^*=0.66 using a logarithmic model.

### Supplementary Table 1: Distribution of *S*_het_ estimates

We provide *S*_het_ estimates in Supplementary Table 1. This file includes the mean *S*_het_ estimates for each gene as well as the upper and lower 95% credibility intervals for each gene estimate. Credibility intervals have precision of 10^−3^ where *S*_het_ > 0.005 and 10^−5^ otherwise.

### Supplementary Table 2: Predicted mode of inheritance for each gene

For each gene, we generate a probability of mode of inheritance (either autosomal dominant or autosomal recessive). Estimates are generated using a logistic regression, trained on the full set of labeled case examples from two clinical exome sequencing programs (Baylor and UCLA)^22,23^. These estimates are applicable for interpretation of genes in cases that are similarly ascertained as these two clinical exome sequencing programs.

### Supplementary Table 3: Analysis of *de novo* variants observed in patients with autism spectrum disorders (ASD)

We use a set of 2,939 ASD case trios and 1,429 control trios with at least one *de novo* variant, which were ascertained from three previous studies^19^. Protein coding *de novo* variants were included for analysis if they did not have a dbSNP rsID and did not appear in any individual in ExAC. Individuals with greater than two *de novo* variants in the coding exome were excluded to avoid potential false positives. Using the Mann-Whitney test, we find that *S*_het_ values are significantly higher in cases versus controls for all coding variants, putative PTVs, putative PTVs with in-frame deletions and insertions, and missense variants that were predicted by PolyPhen-2 to be probably or possibly damaging. As expected, there is no significant difference in *S*_het_ values between ASD cases and controls for synonymous variants or missense variants predicted by PolyPhen-2 to be benign.

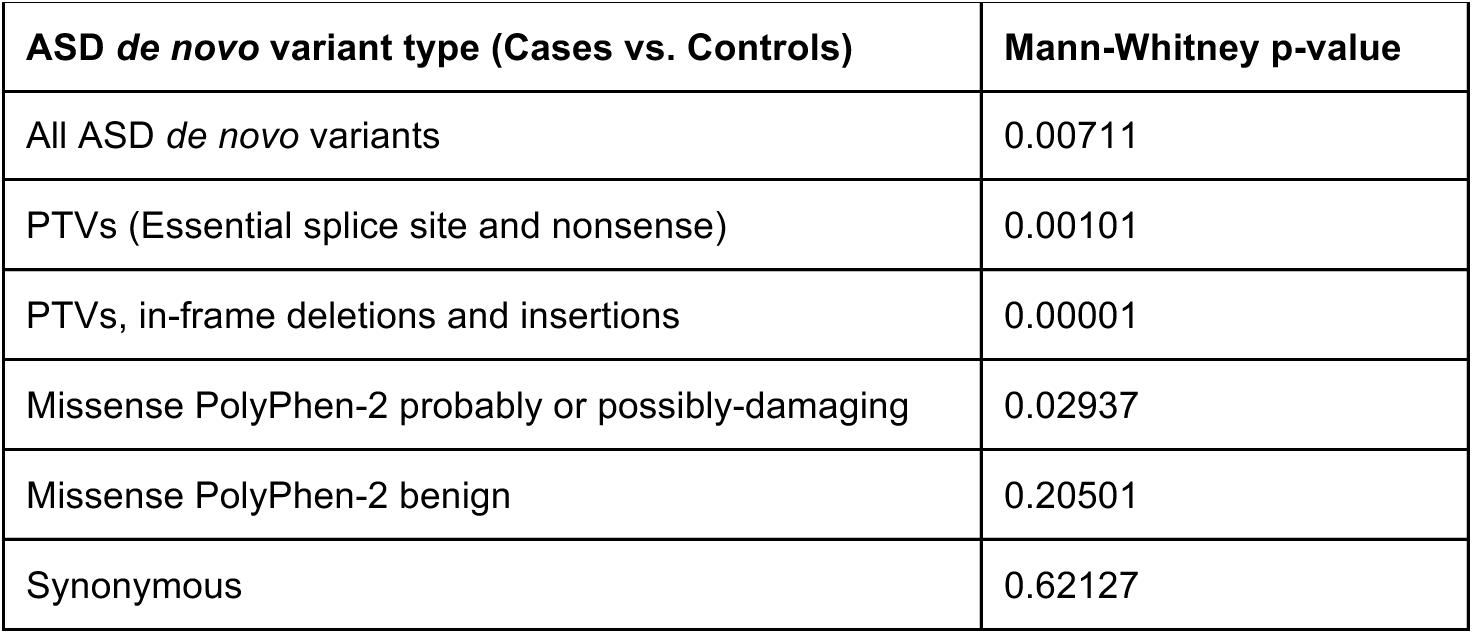

| ASD *de novo* variant type (Cases vs. Controls) | Mann-Whitney p-value |
| --- | --- |
| All ASD *de novo* variants | 0.00711 |
| PTVs (Essential splice site and nonsense) | 0.00101 |
| PTVs, in-frame deletions and insertions | 0.00001 |
| Missense PolyPhen-2 probably or possibly-damaging | 0.02937 |
| Missense PolyPhen-2 benign | 0.20501 |
| Synonymous | 0.62127 |

### Supplementary Table 4: Most published and least published genes from top *S*_het_ decile

Full annotations for the PubMed Score in the top *S*_het_ decile for the top 250 and bottom 250 PubMed genes scores. From the set of genes under the strongest selection (top 10% of *S*_het_ values), we create two sets of 250 genes. We then annotated these lists with the results from neutrally-ascertained screens of gene importance and gene essentiality. We summarize these screens using a heuristic score.

### Supplementary Table 5: Functional analysis terms from DAVID

We include the results of GO term enrichment screening from DAVID that reach Bonferroni corrected significance in genes with *S*_het_ > 0.15, *S*_het_ > 0.25 and *S*_het_ > 0.5.

### Supplementary Table 6: Functional analysis clusters from DAVID

We include the results of functional cluster enrichment screening from DAVID that reach Bonferroni corrected significance in genes with *S*_het_ > 0.15, *S*_het_ > 0.25 and *S*_het_ > 0.5.

## References

1. Mukai, T., Chigusa, S. I., Mettler, L. E. & Crow, J. F. Mutation rate and dominance of genes affecting viability in Drosophila melanogaster. Genetics 72, 335–55 (1972).

2. Deng, H. W. & Lynch, M. Estimation of deleterious-mutation parameters in natural populations. Genetics 144, 349–360 (1996).

3. Wang, T. et al. Identification and characterization of essential genes in the human genome. Science (80-.). 350, 1096–1101 (2015).

4. Williamson, S. H. et al. Simultaneous inference of selection and population growth from patterns of variation in the human genome. Proc. Natl. Acad. Sci. U. S. A. 102, 7882–7 (2005).

5. Boyko, A. R. et al. Assessing the evolutionary impact of amino acid mutations in the human genome. PLoS Genet 4, e1000083(2008).

6. Kryukov, G. V, Pennacchio, L. A. & Sunyaev, S. R. Most rare missense alleles are deleterious in humans: implications for complex disease and association studies. Am. J. Hum. Genet. 80, 727–39 (2007).

7. Kryukov, G. V, Shpunt, A., Stamatoyannopoulos, J. A. & Sunyaev, S. R. Power of deep, all-exon resequencing for discovery of human trait genes. Proc Natl Acad Sci U S A 106, 3871–3876 (2009).

8. Eyre-Walker, A. & Keightley, P. D. The distribution of fitness effects of new mutations. Nat. Rev. Genet. 8, 610–8 (2007).

9. Boyko, A. R. et al. Assessing the evolutionary impact of amino acid mutations in the human genome. PLoS Genet. 4, e1000083 (2008).

10. Do, R. et al. No evidence that selection has been less effective at removing deleterious mutations in Europeans than in Africans. Nat. Genet. 47, 126–131 (2015).

11. Fu, W., Gittelman, R. M., Bamshad, M. J. & Akey, J. M. Characteristics of neutral and deleterious protein-coding variation among individuals and populations. Am. J. Hum. Genet. 95, 421–36 (2014).

12. Lohmueller, K. E. The distribution of deleterious genetic variation in human populations. Curr. Opin. Genet. Dev. 29, 139–46 (2014).

13. Gravel, S. When Is Selection Effective? Genetics 203, 451–62 (2016).

14. Williamson, S., Fledel-Alon, A. & Bustamante, C. D. Population genetics of polymorphism and divergence for diploid selection models with arbitrary dominance. Genetics 168, 463–75 (2004).

15. Balick, D. J., Do, R., Cassa, C. A., Reich, D. & Sunyaev, S. R. Dominance of Deleterious Alleles Controls the Response to a Population Bottleneck. PLoS Genet. 11, e1005436 (2015).

16. Simons, Y. B., Turchin, M. C., Pritchard, J. K. & Sella, G. The deleterious mutation load is insensitive to recent population history. Nat. Genet. 46, 220–224 (2014).

17. Lek, M. et al. Analysis of protein-coding genetic variation in 60, 706 humans. Nature 536, 285–291 (2016).

18. MacArthur, D. G. et al. A systematic survey of loss-of-function variants in human protein-coding genes. Science (80-.). 335, 823–828 (2012).

19. Samocha, K. E. et al. A framework for the interpretation of de novo mutation in human disease. Nat. Genet. 46, 944–50 (2014).

20. Francioli, L. C. et al. Genome-wide patterns and properties of de novo mutations in humans. Nat. Genet. 47, 822–6 (2015).

21. Solomon, B. D., Nguyen, A.-D., Bear, K. A. & Wolfsberg, T. G. Clinical genomic database. Proc. Natl. Acad. Sci. U. S. A. 110, 9851–5 (2013).

22. Yang, Y. et al. Molecular Findings Among Patients Referred for Clinical Whole-Exome Sequencing. JAMA (2014). doi:10.1001/jama.2014.14601

23. Lee, H. et al. Clinical Exome Sequencing for Genetic Identification of Rare Mendelian Disorders. JAMA (2014). doi:10.1001/jama.2014.14604

24. Saleheen, D. et al. Human knockouts in a cohort with a high rate of consanguinity. (2015). doi:10.1101/031518

25. Koscielny, G. et al. The International Mouse Phenotyping Consortium Web Portal, a unified point of access for knockout mice and related phenotyping data. Nucleic Acids Res. 42, D802–9 (2014).

26. Georgi, B., Voight, B. F. & Bućan, M. From Mouse to Human: Evolutionary Genomics Analysis of Human Orthologs of Essential Genes. PLoS Genet. 9, e1003484 (2013).

27. Roessler, E. et al. Mutations in the human Sonic Hedgehog gene cause holoprosencephaly. Nat. Genet. 14, 357–60 (1996).

28. Kang, S., Graham, J. M., Olney, A. H. & Biesecker, L. G. GLI3 frameshift mutations cause autosomal dominant Pallister-Hall syndrome. Nat. Genet. 15, 266–8 (1997).

29. Vortkamp, A., Gessler, M. & Grzeschik, K. H. GLI3 zinc-finger gene interrupted by translocations in Greig syndrome families. Nature 352, 539–40 (1991).

30. Wild, A. et al. Point mutations in human GLI3 cause Greig syndrome. Hum. Mol. Genet. 6, 1979–84 (1997).

31. Roessler, E. et al. Loss-of-function mutations in the human GLI2 gene are associated with pituitary anomalies and holoprosencephaly-like features. Proc. Natl. Acad. Sci. U. S. A. 100, 13424–9 (2003).

32. Chiang, C. et al. Cyclopia and defective axial patterning in mice lacking Sonic hedgehog gene function. Nature 383, 407–13 (1996).

33. Hui, C. C. & Joyner, A. L. A mouse model of greig cephalopolysyndactyly syndrome: the extra-toesJ mutation contains an intragenic deletion of the Gli3 gene. Nat. Genet. 3, 241–6 (1993).

34. Mo, R. et al. Specific and redundant functions of Gli2 and Gli3 zinc finger genes in skeletal patterning and development. Development 124, 113–23 (1997).

35. Huang, D. W., Sherman, B. T. & Lempicki, R. A. Systematic and integrative analysis of large gene lists using DAVID bioinformatics resources. Nat. Protoc. 4, 44–57 (2008).

36. Seidman, J. G. & Seidman, C. Transcription factor haploinsufficiency: when half a loaf is not enough. J. Clin. Invest. 109, 451–455 (2002).

37. Huttlin, E. L. et al. The BioPlex Network: A Systematic Exploration of the Human Interactome. Cell 162, 425–40 (2015).

38. NCBI Resource Coordinators. Database resources of the National Center for Biotechnology Information. Nucleic Acids Res. 41, D8–D20 (2013).

39. Raychaudhuri, S. et al. Identifying relationships among genomic disease regions: Predicting genes at pathogenic SNP associations and rare deletions. PLoS Genet. 5, (2009).

40. Agrawal, A. F. & Whitlock, M. C. Inferences about the distribution of dominance drawn from yeast gene knockout data. Genetics 187, 553–566 (2011).

41. Simmons, M. J. & Crow, J. F. Mutations Affecting Fitness in Drosophila Populations. Annu. Rev. Genet. 11, 49–78 (1977).

42. Wright, S. Evolution in Mendelian populations. 1931. Bull. Math. Biol. 52, 241–95–7 (1990).

43. Petrovski, S. et al. The Intolerance of Regulatory Sequence to Genetic Variation Predicts Gene Dosage Sensitivity. PLoS Genet. 11, e1005492 (2015).

44. Kiezun, A. et al. Exome sequencing and the genetic basis of complex traits. Nat. Genet. 44, 623–30 (2012).

45. Messer, P. W. SLiM: simulating evolution with selection and linkage. Genetics 194, 1037–9 (2013).

46. Tennessen, J. A. et al. Evolution and Functional Impact of Rare Coding Variation from Deep Sequencing of Human Exomes. Science (80-.). 337, 64–69 (2012).

47. Karczewski, K. LOFTEE (Loss-Of-Function Transcript Effect Estimator). (2015). at <https://github.com/konradjk/loftee>

48. Ayadi, A. et al. Mouse large-scale phenotyping initiatives: overview of the European Mouse Disease Clinic (EUMODIC) and of the Wellcome Trust Sanger Institute Mouse Genetics Project. Mamm. Genome 23, 600–10 (2012).

49. McKusick-Nathans Institute of Genetic Medicine MD), J. H. U. (Baltimore. Online Mendelian Inheritance in Man, OMIM®. at <http://omim.org/>

50. Cassa, C. A., Tong, M. Y. & Jordan, D. M. Large numbers of genetic variants considered to be pathogenic are common in asymptomatic individuals. Hum Mutat 34, 1216–1220 (2013).

51. Tong, M. Y., Cassa, C. A. & Kohane, I. S. Automated validation of genetic variants from large databases: ensuring that variant references refer to the same genomic locations. Bioinformatics 27, 891–893 (2011).

52. Zhang, J. et al. Germline Mutations in Predisposition Genes in Pediatric Cancer. N. Engl. J. Med. 373, 2336–2346 (2015).

